# VINCIT: A virus-incorporated biomaterial niche enables spatially controlled in vivo CAR T cell engineering

**DOI:** 10.64898/2026.11.16.767176

**Authors:** Kailei D. Goodier, Yining Zhu, Christine Wei, Zhuoqing Jiang, Yili Jiang, Xiaoya Lu, Sabrina Chen, Autumn H. Greco, Di Yu, Xiang Liu, Joseph Choy, Yunhe Su, Jingyao Ma, Jinghan Lin, Jiayuan Kong, Evan Wang, Dylan Luong, D. Scott Wilson, Joshua C. Doloff, Jonathan P. Schneck, Maximilian F. Konig, Hai-Quan Mao

**Affiliations:** Institute for NanoBioTechnology, Johns Hopkins University, Baltimore, MD 21218, USA; Department of Materials Science and Engineering, Johns Hopkins University, Baltimore, MD 21218, USA; Translational Therapeutics & Regenerative Engineering Center, Johns Hopkins University School of Medicine, Baltimore, MD 21231, USA; Department of Biomedical Engineering, Johns Hopkins University School of Medicine, Baltimore, MD 21218, USA; Department of Chemistry, Johns Hopkins University, Baltimore, MD 21218, USA; Institute for Cell Engineering, Johns Hopkins University School of Medicine, Baltimore, MD 21231, USA; Department of Pathology, Johns Hopkins University School of Medicine, Baltimore, MD 21231, USA; Department of Medicine, Johns Hopkins University School of Medicine, Baltimore, MD 21231, USA; Sidney-Kimmel Comprehensive Cancer Center, Johns Hopkins University, Baltimore, MD, USA; Bloomberg-Kimmel Institute for Cancer Immunotherapy, Johns Hopkins University, Baltimore, MD, USA; Department of Oncology, Johns Hopkins University School of Medicine, Baltimore, MD 21231, USA; Translational Immunoengineering Center, Johns Hopkins University School of Medicine, Baltimore, MD 21231, USA; Division of Rheumatology, Department of Medicine, Johns Hopkins University School of Medicine, Baltimore, MD 21224, USA; Center for Autoimmunity and Immuno-Oncology, Johns Hopkins University School of Medicine, Baltimore, MD 21287, USA

## Abstract

In vivo generation of chimeric antigen receptor (CAR) T cells could bypass the complex manufacturing required for conventional cell therapies, but systemic gene delivery remains limited by off-target exposure and inefficient access to target immune cells. Here, we develop VINCIT (Virus-Incorporated Niche for Cellular ImmunoTherapy), an injectable nanofiber–hydrogel composite that establishes a localized immune-engineering niche for CAR T cell generation in vivo. VINCIT spatially retains viral vectors while recruiting endogenous immune cells, enabling local transduction of T cells, after which the engineered T cells enter the circulation. VINCIT spatially confines transgene expression compared with intravenous viral delivery and markedly enhances T cell transduction compared with subcutaneously injected free virus. VINCIT is compatible with both adeno-associated and lentiviral vectors, and CAR T cells generated in vivo exhibit antigen-specific effector responses and mediate CD19^+^ B cell depletion. In a humanized mouse model of lymphoma, a single VINCIT administration produces sustained tumor control and extends median survival from 32 to 110 days. Mechanistically, efficient CAR T cell generation requires coordinated interactions among multiple immune populations, with neutrophil depletion markedly reducing cellular recruitment and subsequent T cell programming. These findings establish a localized biomaterial niche for spatially controlled in vivo immune cell engineering.

## INTRODUCTION

Chimeric antigen receptor (CAR) T cell therapies have reshaped the treatment landscape for B cell malignancies^1,2^. Autologous CD19-targeted cellular therapies, including tisagenlecleucel and axicabtagene ciloleucel, can induce durable complete remissions in patients with relapsed or refractory leukemia and lymphoma^3–6^. This clinical success has driven rapid expansion into other hematologic cancers, solid tumors, and refractory autoimmune diseases, where early results, including drug-free remissions in systemic lupus erythematosus, highlight the broader therapeutic potential of engineered T cells^6–8^. Despite these advances, the clinical impact of CAR T therapy remains constrained by accessibility, manufacturing complexity, and cost. Current autologous products require lymphodepleting chemotherapy to enable robust engraftment, exposing patients to immunosuppression, infection risk, and treatment-related morbidity^9–11^. Manufacturing timelines spanning weeks from leukapheresis to infusion can exceed the time available for patients with rapidly progressive disease, contributing to substantial attrition before treatment delivery^12^. Together, these challenges establish ex vivo manufacturing as a central bottleneck limiting the scalability and timely deployment of CAR T therapies.

Considerable effort has therefore focused on bypassing ex vivo CAR T manufacturing^13^. Allogeneic off-the-shelf products derived from healthy donors offer immediate availability but require extensive genome editing to prevent graft-versus-host disease, and remain limited by host immune rejection and limited persistence^14,15^. In vivo T cell programming has emerged as an alternative to ex vivo CAR T manufacturing. Targeted lipid nanoparticles delivering CAR mRNA have achieved transient T cell reprogramming and B cell depletion in non-human primates, with multiple candidates in early-stage trials. Still, mRNA-based approaches yield only transient expression, which limits therapeutic potency. Viral vectors offer the potential for durable in vivo T cell engineering, but systemic administration creates a tradeoff between sufficient vector exposure for efficient transduction and systemic or off-target effects^16,17^. These limitations motivate strategies that spatially confine vector exposure while maintaining access to target immune cells^18–20^.

Biomaterial scaffolds have shown that spatially restricted niches can support T cell activation, transduction, and expansion while reducing systemic exposure^21,22^. Implantable alginate scaffolds have enabled rapid CAR transduction of pre-loaded immune cells for tumor control in xenograft models of lymphoma^23^. These scaffold-based approaches, however, rely on patient cell isolation and ex vivo loading before implantation, preserving a critical ex vivo processing step^23,24^ and limiting scalability. Thus, current biomaterial strategies improve spatial control but do not bypass the dependence on ex vivo cell manipulation.

We previously developed a biostimulatory nanofiber-hydrogel composite (NHC) that mimics the microarchitecture and mechanical properties of soft-tissue extracellular matrix^25^. HA was chosen because it is a native matrix glycosaminoglycan forming a hydrated, cell-permissive network, while covalently conjugated PCL nanofibers provide structural reinforcement, decoupling stiffness from the porosity needed for host cell infiltration. This system recruits host immune cells following subcutaneous injection, and has demonstrated favorable biocompatibility and integration with host tissue. These properties suggest the potential of establishing a localized immune-engineering niche capable of recruiting endogenous immune cells and coordinating their genetic programming in situ. An approach that integrates in vivo cell recruitment, spatially confined gene delivery, and efficient T cell programming within a single scaffold remains lacking.

Here, we introduce a Virus-Incorporated Niche for Cellular ImmunoTherapy (VINCIT) that enables in vivo CAR T cell generation through a single subcutaneous injection, without ex vivo cell processing or loading. VINCIT integrates recruitment of endogenous immune cells with spatially confined viral delivery, enabling recruited T cells to be transduced in situ and subsequently enter the systemic circulation. We evaluate this strategy using both AAV and lentiviral vectors and demonstrate localized viral retention, enhanced T cell transduction, functional CAR T cell generation, and durable tumor control. We further investigate the multicellular immune cell interactions within VINCIT that support efficient in vivo T cell programming.

## RESULTS

### VINCIT localizes viral gene delivery to enable in situ T cell recruitment and transduction

To enable *in vivo* CAR T cell generation without ex vivo cell manipulation, we developed VINCIT by incorporating viral vectors into an NHC that functions as a localized immune-engineering niche (**Fig. 1a**). Throughout, NHC refers to the unloaded nanofiber–hydrogel composite and VINCIT to the same composite loaded with viral vector. NHC was selected based on its established biocompatibility and safety^26^, and its ability to recruit host immune cells following subcutaneous injection. Structurally, NHC consists of a hyaluronic acid (HA) hydrogel network covalently integrated with electrospun poly(ε-caprolactone) (PCL) nanofiber fragments, forming a mechanically stable yet highly porous matrix^27^. This composite architecture preserves an interconnected macroporous network that supports immune cell infiltration and retention, with a shear storage modulus G’ of ∼300 Pa (**Supplementary Fig. 1**). Scanning electron microscopy revealed a well-connected macroporous structure with average pore sizes of approximately 50 μm (**Supplementary Fig. 2**), closely resembling the microarchitecture of soft-tissue extracellular matrix and enabling efficient host immune cell recruitment^28–30^.

**Figure 1.**
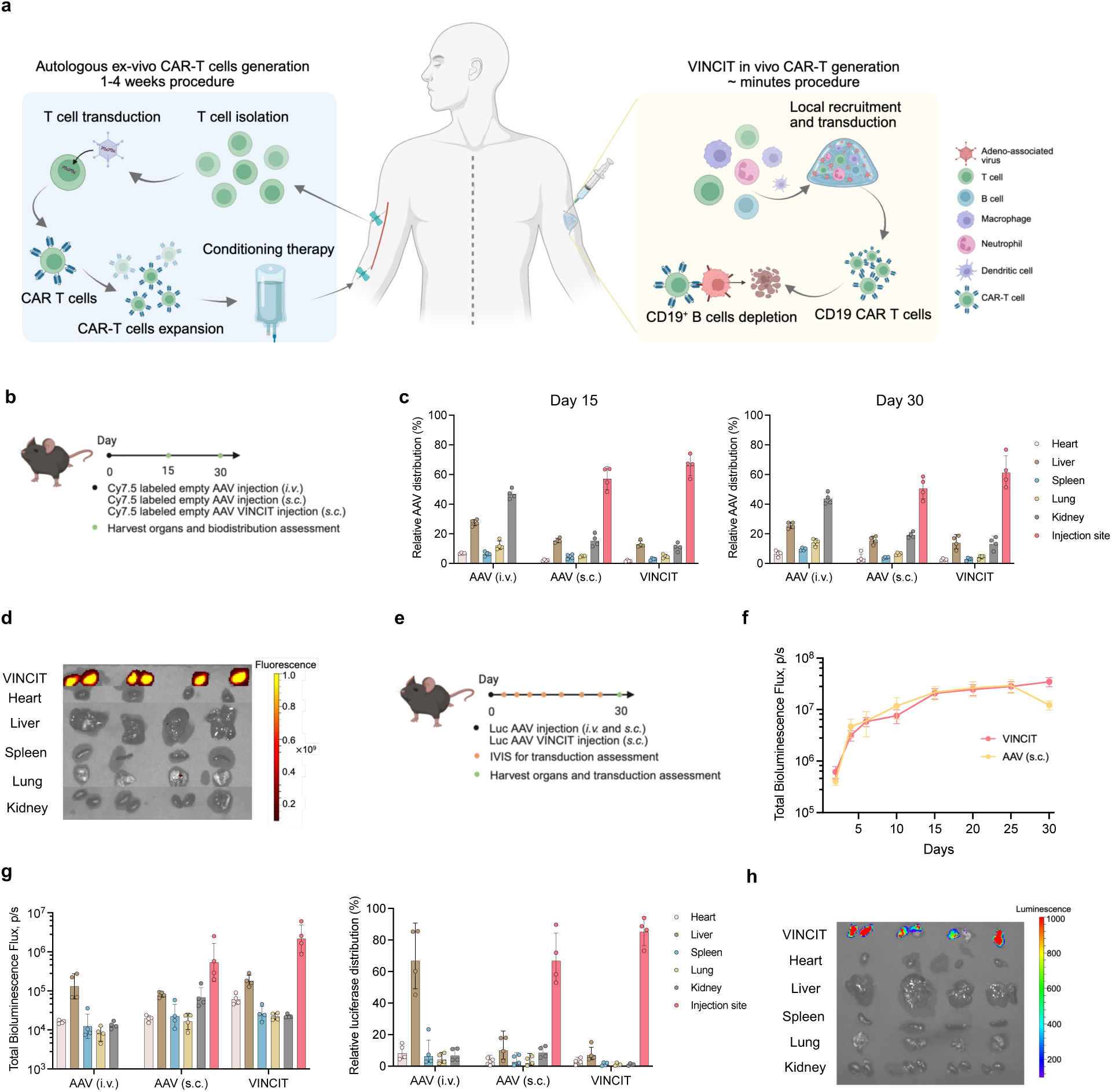
VINCIT enables localized viral gene delivery for *in vivo* CAR T cell engineering. **a.** Schematic comparing conventional CAR T cell manufacturing with VINCIT-mediated *in vivo* CAR T cell generation. Autologous CAR T production (left) is a multi-step process requiring leukapheresis, T cell isolation, ex vivo transduction, and expansion over 1-4 weeks before infusion into patients. In contrast, VINCIT (right) enables *in vivo* CAR T generation through a single subcutaneous injection, where the nanofiber-hydrogel composite serves as a transient immunological niche that recruits host T cells, transduces them locally with CAR-encoding viral vectors, after which the engineered CAR T cells enter the circulation to mediate target cell depletion. **b**–**d,** C57BL/6 mice received Cy7.5-labeled empty AAV via intravenous injection (*i.v.*), subcutaneous injection (*s.c.*), or subcutaneous injection with VINCIT. Organs were harvested at days 15 and 30 for assessment. Experimental timeline (**b**) and quantification of AAV distribution following different administration routes on days 15 and 30 (**c**). *Ex vivo* fluorescence imaging of harvested organs following Cy7.5-labeled AAV VINCIT delivery at day 15. **e**–**h**, Transduction localization assessed by bioluminescence imaging (IVIS). **e**, Experimental timeline. Mice received Luc AAV via the indicated routes and were monitored by IVIS imaging. **f**, Luciferase expression (IVIS bioluminescence signal) at the injection site over 30 days. **g,** Total bioluminescence and relative distribution of luciferase expression among different organs at day 30 across different delivery groups. **h,** Bioluminescence imaging of harvested organs following Luc AAV VINCIT delivery. Data are from n = 4 biologically independent samples. Data represent mean ± s.e.m. Panels **a, b,** and **e** were prepared in BioRender.

We first evaluated whether VINCIT could achieve localized viral retention compared to conventional delivery routes. Mice received Cy7.5-labeled AAV via three routes: intravenous injection, subcutaneous injection of free virus, or subcutaneous injection of virus premixed with the NHC, forming VINCIT (**Fig. 1b**). Organs were harvested at days 15 and 30 to assess biodistribution. Intravenous administration resulted in predominant hepatic and renal accumulation. In contrast, VINCIT increased local viral retention at the injection site, with over 60% of the total signal at the injection site at both time points, and correspondingly reduced distribution to off-target organs (**Fig. 1c, d**; **Supplementary Figs. 3** and **4**).

We next examined whether the virus confined within VINCIT remained functionally active for cellular transduction over time. Using an AAV encoding luciferase, we monitored bioluminescence signal using the in vivo imaging system (IVIS) over 30 days (**Fig. 1e**). Intravenous delivery of AAV produced liver-dominant transduction, with approximately 70% of total signal localized to the liver, consistent with off-target hepatic gene expression. Subcutaneous (s.c.) injection of free AAV resulted in ∼68% of luciferase expression signal localized at the injection site (*p* < 0.01, vs. i.v. AAV group), accompanied by the corresponding systemic spread (∼12% in the liver and ∼9% in the kidney). VINCIT confined more than 85% of the bioluminescent signal to the injection site (*p* < 0.01, vs. i.v. AAV group), while maintaining sustained transduction activity, demonstrating more spatially restricted and sustained transgene expression within the scaffold (**Fig. 1g, h**; **Supplementary Figs. 5** and **6**). These results indicate that VINCIT retains viral particles locally and preserves their functional activity for in situ gene transfer. Together, these results demonstrate that VINCIT achieves spatial confinement of viral gene delivery while sustaining functional transduction within a localized tissue niche.

Since NHC can recruit host immune cells and retain them within the niche, and VINCIT spatially confines AAV delivery at the injection site, we next examined whether it supports in situ transduction of the recruited cells. More importantly, subcutaneously delivered free AAV mainly transduces local tissue rather than immune cells. In mice, lacZ mapping showed the panniculus carnosus, a skeletal muscle layer of the skin, as the primary transduced tissue after s.c. rAAV injection^31^. Although free s.c. AAV efficiently transduced local tissue, only a small fraction of transduced cells were CD3^+^ T cells (**Supplementary Fig. 7**), indicating that localization of viral exposure alone is insufficient for efficient T cell programming. We therefore compared VINCIT with virus-loaded HA hydrogel (ViHA) to isolate the NHC contribution and assess the scaffold’s effect on immune cell recruitment, retention, and transduction efficiency. Mice received s.c. injections of VINCIT or ViHA loaded with a tdTomato-encoding AAV were sacrificed on days 3, 7, and 30 for analysis (**Fig. 2a**).

**Figure 2.**
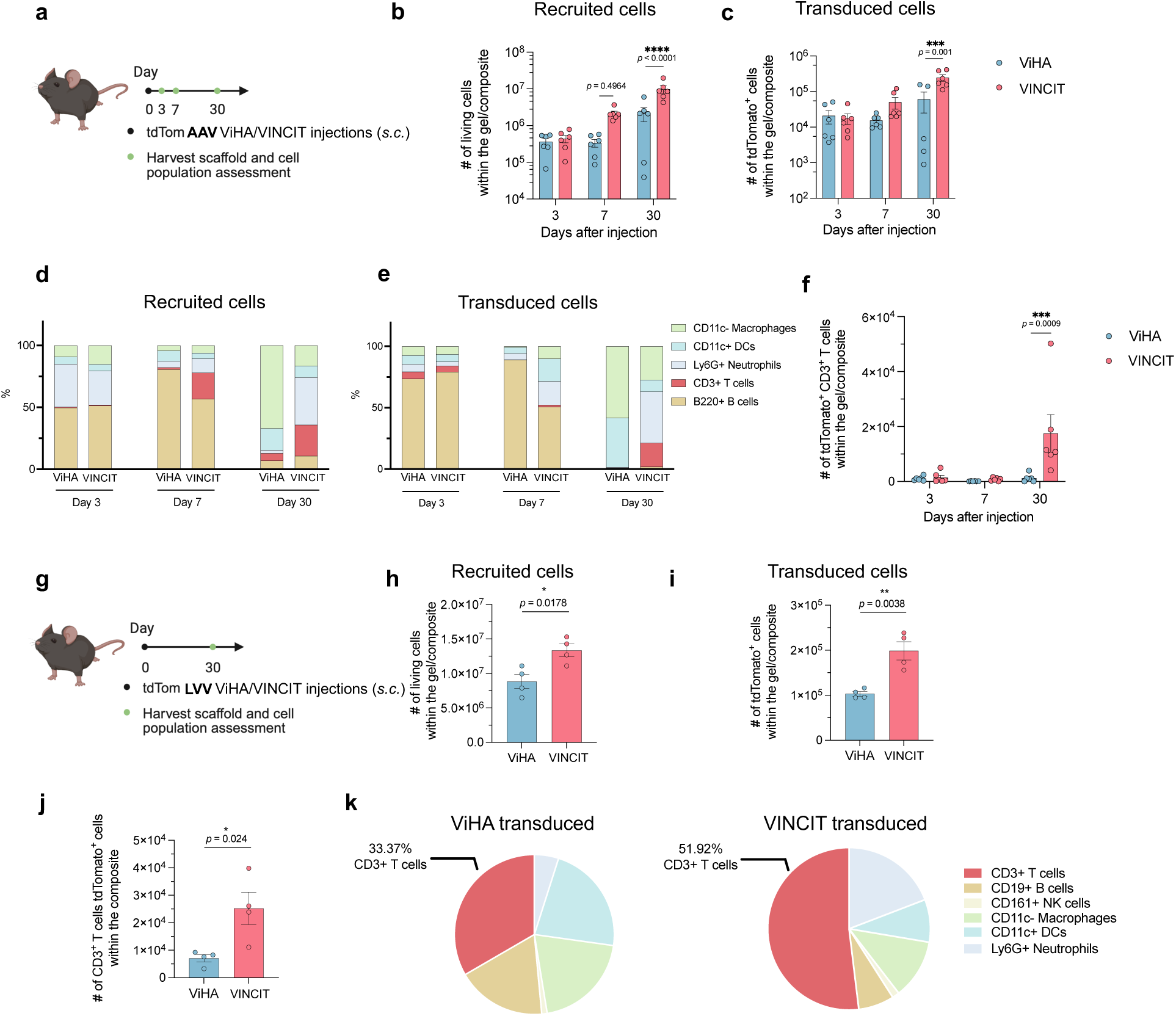
Localized viral delivery by VINCIT drives immune cell recruitment and T cell transduction. **a**–**f**, Cell recruitment and transduction in ViHA and VINCIT with tdTomato AAV. **a**, Experimental timeline. C57BL/6 mice received subcutaneous injection of ViHA or VINCIT with tdTomato AAV. Scaffolds were harvested at days 3, 7, and 30 for analysis. **b**, Total number of recruited cells within ViHA and VINCIT over time. **c**, Total number of transduced cells within ViHA and VINCIT over time. **d, e**, Composition of recruited cells (d) and transduced cells (e) in ViHA and VINCIT at days 3, 7, and 30, showing distribution of macrophages (CD11c^−^), dendritic cells (CD11c^+^), neutrophils (Ly6G^+^), CD3^+^ T cells, and B220^+^ B cells. **f**, Quantification of transduced CD3^+^ T cells within ViHA and VINCIT over time. **g**–**k**, Cell recruitment and transduction in ViHA and VINCIT using lentiviral vectors (LVV). **g**, Experimental timeline. Mice received subcutaneous injection of ViHA or VINCIT with tdTomato LVV and were assessed at day 30. **h**, Total number of recruited cells in ViHA and VINCIT at day 30. **i**, Total number of transduced cells. **j**, Number of transduced CD3^+^ T cells. **k**, Composition of transduced cells in ViHA and VINCIT. Data are from n = 6 (**b**–**f**) or 4 (**h**–**k**) biologically independent samples. Data were analyzed using an unpaired t-test. NS: *p* > 0.05, \**p* < 0.05, \*\**p* < 0.01, \*\*\**p* < 0.001, \*\*\*\**p* < 0.0001. Panels **a** and **g** were created in BioRender.

Both scaffolds recruited host cells progressively; however, VINCIT mediated substantially more cellular infiltration, containing approximately 4-fold more total cells than ViHA by day 30 (**Fig. 2b**). The composition of recruited cells shifted over time (**Fig. 2d**; **Supplementary Fig. 8**). At day 3, both groups were dominated by neutrophils and B cells. By day 7, T cells began to infiltrate, with greater accumulation in VINCIT compared to ViHA. By day 30, VINCIT exhibited a pronounced enrichment of T cells and neutrophils, whereas ViHA remained predominantly macrophage-enriched.

By day 30, VINCIT generated approximately 4-fold more total transduced cells than ViHA (**Fig. 2c**). The composition of transduced cells largely reflected recruited cell composition (**Fig. 2e**), indicating proportional gene transfer across major immune subsets. Notably, VINCIT generated a striking ∼15-fold increase in transduced CD3⁺ T cells compared to ViHA by day 30 (**Fig. 2f**), demonstrating that enhanced T-cell recruitment within VINCIT translates into substantially greater in situ T-cell transduction within the niche.

To assess whether VINCIT is compatible with different viral vector systems, we repeated these experiments using lentiviral vectors (LVV), which enable durable transgene integration and have recently advanced into clinical evaluation for in vivo CAR T cell generation. Consistent with the AAV results, VINCIT recruited significantly more total cells than ViHA at day 30 (**Fig. 2h**), and this enhanced recruitment translated into greater total transduction (**Fig. 2i**) and a higher yield of transduced CD3^+^ T cells (**Fig. 2j, k**). These findings demonstrate that the enhanced recruitment and transduction enabled by VINCIT are not specific to a single vector system, but extend across both non-integrating (AAV) and integrating (LVV) platforms. Together, these results demonstrate that VINCIT establishes a permissive niche that drives immune cell recruitment and enhances in situ T cell transduction for both AAV and LVV.

### VINCIT-generated CAR T cells circulate systemically and exhibit functional cytotoxicity

Having established that VINCIT recruits and transduces T cells in situ, we next asked whether these engineered cells exit the local niche and behave as circulating therapeutic T cells. To assess systemic trafficking, mice received subcutaneous injections of VINCIT or ViHA loaded with a tdTomato-encoding AAV, and spleens were harvested at days 15 and 30 for analysis (**Fig. 3a**).

**Figure 3.**
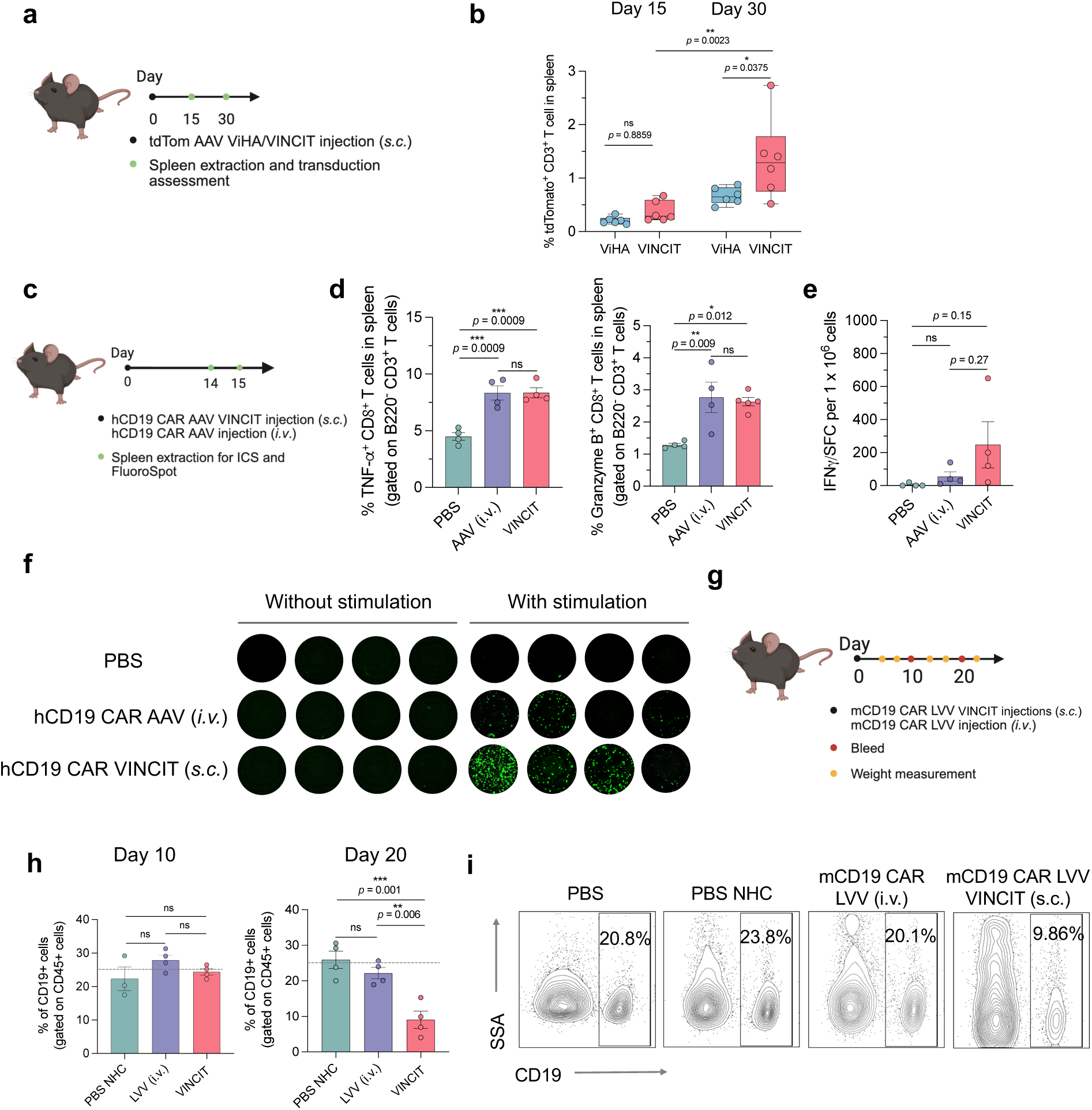
*In vivo* CAR T cells generated by VINCIT are functional and mediate B cell depletion. **a, b**, Cell recruitment and transduction in ViHA and VINCIT using tdTomato AAV. **a**, Experimental timeline. C57BL/6 mice received subcutaneous injection of ViHA or VINCIT. Spleens were harvested at days 15 and 30 for analysis. **b**, Percentage of transduced CD3^+^ T cells among total live cells at days 15 and 30. **c**–**f**, Functional assessment of VINCIT-generated CAR T cells. **c**, Experimental timeline. C57BL/6 mice received hCD19 CAR AAV via intravenous injection (i.v.) or subcutaneous VINCIT injection. Splenocytes were harvested at day 15 for intracellular cytokine staining (ICS) and FluoroSpot analysis. **d–f**, Splenocytes were stimulated ex vivo with hCD19+ lysate. **d**, Quantification of TNF-α and Granzyme B production by splenocytes following ex vivo stimulation. **e**, Quantification of IFN-γ–producing cells by FluoroSpot assay. **f**, IFN-γ FluoroSpot images showing IFN-γ–producing cells from individual mice in each group. **g**–**i**, B cell depletion by CAR T cells assessed in different groups. **g**, Experimental timeline. Mice received mCD19 CAR LVV via i.v. injection or s.c. VINCIT injection. Peripheral blood was collected on days 10 and 20. **h**, Quantification of CD19^+^ B cells in peripheral blood at days 10 and 20. The dashed line indicates the baseline B cell level in PBS-treated mice. **i**, Representative flow cytometry plots of CD19^+^ B cells. Data are from n = 6 (**b**) or 4 (**d**–**i**) biologically independent samples. Data were analyzed using an unpaired t-test for **b**, and one-way ANOVA and Tukey’s multiple comparisons test for **d, e, h**. NS: *p* > 0.05, \**p* < 0.05, \*\**p* < 0.01, \*\*\**p* < 0.001, \*\*\*\**p* < 0.0001. Panels **a, c, g** were created in BioRender.

On day 15, only a small population of transduced T cells was detectable in the spleen, with no significant difference between VINCIT- and ViHA-treated groups. By day 30, transduced T cells comprised approximately 1.5% of the splenocyte population in VINCIT-treated mice, significantly higher than in the ViHA-treated control group (<1%, p < 0.05) (**Fig. 3b**). This increase over time is consistent with peripheral trafficking of locally transduced T cells from the VINCIT niche.

We next assessed whether VINCIT generates functionally competent CAR T cells *in vivo*. VINCIT matrices were loaded with an AAV encoding anti-human CD19 CAR and injected subcutaneously into C57BL/6 mice (**Fig. 3c**). For comparison, a separate cohort received the same AAV dose via i.v. administration (**Fig. 3c**). Splenocytes were harvested on day 15 for functional evaluation (**Fig. 3d**). Upon ex vivo stimulation with lysate derived from hCD19⁺ Daudi cells, CD8⁺ T cells from VINCIT-treated mice exhibited ∼2-fold higher expression of TNF-α and granzyme B compared with the PBS control, reaching levels comparable to those observed following systemic AAV delivery (**Fig. 3e**). Consistent with these findings, IFN-γ FluoroSpot analysis revealed a marked increase in antigen-responsive cells in VINCIT-treated mice, with ∼50-fold more IFN-γ–secreting cells than the PBS control and ∼4.5-fold more than mice receiving intravenous AAV (**Fig. 3f, g**). These data demonstrate that VINCIT-generated T cells exhibit antigen-specific activation and effector cytokine production, consistent with functional CAR T cell responses.

To determine whether VINCIT-generated CAR T cells exhibit functional activity *in vivo*, we next assessed CD19⁺ B cell depletion as a pharmacodynamic readout. B cell aplasia is a well-established surrogate marker of CD19 CAR T cell activity and is routinely used clinically to monitor therapeutic persistence and function^32^. Because this readout requires depletion of endogenous murine B cells, these experiments used a CAR directed against murine CD19, which was available to us in a lentiviral backbone. Mice were treated with this LVV encoding a murine CD19-targeted CAR, delivered either via VINCIT or through i.v. administration, with the virus free NHC and PBS injections as controls (**Fig. 3g**). Peripheral blood analysis revealed substantial B cell depletion in VINCIT-treated mice by day 20, with CD19⁺ B cell frequencies reduced to approximately 8%, representing a 65% reduction relative to the PBS control group. In contrast, mice receiving i.v. CAR LVV retained approximately 22% CD19⁺ B cells, comparable to the control groups (**Fig. 3h, i**). Together, these results demonstrate that VINCIT generates systemically circulating, antigen-responsive CAR T cells that exhibit robust effector function and mediate effective target cell elimination *in vivo*.

### VINCIT-mediated in vivo CAR T generation suppresses tumor growth and prolongs survival

To evaluate VINCIT’s therapeutic efficacy, we used a humanized mouse model of aggressive CD19⁺ B cell lymphoma. NSG mice were injected intravenously with 1 × 10^6^ luciferase-expressing Daudi B cells on day 0. To reconstitute a human immune compartment, mice received an intravenous injection of 1 × 10^7^ human PBMCs on day 5. On day 7, animals received a single dose of human CD19 CAR-encoding AAV delivered either subcutaneously via VINCIT or systemically via intravenous injection. Control groups received PBS (no treatment) or virus free NHC (**Fig. 4a**). Tumor progression was monitored longitudinally by IVIS. This model enables simultaneous evaluation of *in vivo* CAR T generation, systemic trafficking, and antitumor efficacy.

**Figure 4.**
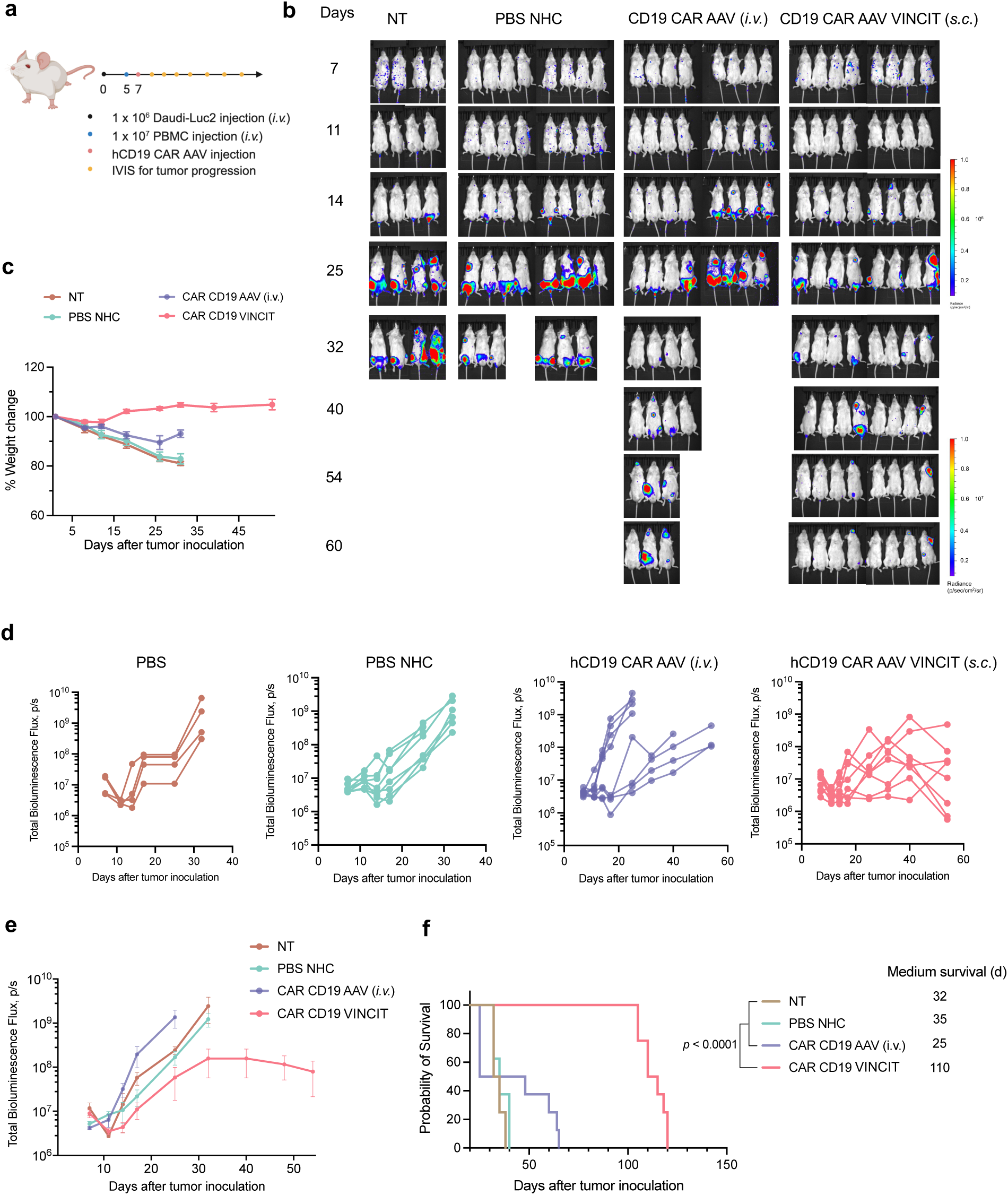
VINCIT controls tumor growth and extends survival in a humanized lymphoma model. **a**, Experimental timeline. NSG mice received intravenous injection of Daudi-Luc2 cells on day 0 and human PBMCs on day 5 to establish a humanized model of B cell lymphoma. On day 7, mice (n = 4 for PBS groups and n = 8 for other groups) were treated with PBS, hCD19 CAR AAV via intravenous injection (*i.v.*), virus free NHC, hCD19 CAR AAV VINCIT via subcutaneous injection (*s.c.*). Tumor progression was monitored by IVIS imaging. **b**, Bioluminescence images of tumor-bearing mice at the indicated time points. **c**, Body weight changes over time. **d**, Individual tumor burden curves for each treatment group measured by total bioluminescence flux. **e**, Average bioluminescence flux over time across treatment groups. **f**, Survival curves for animals treated with indicated conditions. Data are from n = 8 biologically independent samples. Significance in **f** was calculated using a log-rank Mantel-Cox test. \*\*\*\**p* < 0.0001. Panel **a** was created in BioRender.

Control mice (PBS or NHC only) exhibited rapid tumor progression, reaching high tumor burden by day 25 (**Fig. 4b**). In contrast, VINCIT-treated mice displayed sustained tumor suppression, with markedly reduced tumor burden compared with all other treatment groups throughout the 60-day observation period (**Fig. 4d, e, Supplementary Fig. 9**). Notably, intravenous delivery of the same CAR-encoding AAV failed to achieve comparable tumor control, underscoring the therapeutic advantage of localized gene delivery in this model. VINCIT-treated mice maintained stable body weight throughout the observation period, whereas the PBS, NHC, and intravenous CAR AAV groups showed progressive weight loss that paralleled their increasing tumor burden (Fig. 4c). Body weight was therefore preserved only in the group that controlled tumor growth, indicating good tolerability of VINCIT. Consistent with these findings, survival analysis demonstrated a marked therapeutic benefit in the VINCIT cohort (**Fig. 4f**). Untreated mice had a median survival of 32 days, while mice receiving the NHC alone exhibited a median survival of 35 days, confirming that the scaffold itself did not confer antitumor activity. By comparison, VINCIT-treated mice achieved a median survival of 110 days. These results demonstrate that a single administration of VINCIT is sufficient to produce sustained therapeutic benefit in a human B cell lymphoma model.

To assess the systemic inflammatory response to VINCIT, we measured serum cytokines three days after treatment with empty AAV. VINCIT did not induce detectable elevations in IL-6, TNF-α, IL-10, IL-12, or IFN-γ compared with PBS-, free AAV-, or ViHA-treated controls (**Supplementary Fig. 10**). These findings indicate that localized VINCIT delivery was not associated with detectable systemic cytokine elevation under the conditions tested. Together, these results demonstrate that localized *in vivo* CAR T cell generation via VINCIT enables durable tumor control and substantial survival benefit following a single injection.

### VINCIT establishes a coordinated immune niche that enables in vivo CAR T cell generation

We next explored the immune interactions underlying VINCIT-mediated efficacy and whether VINCIT functions as an active immune niche rather than solely as a localized gene-delivery depot. We first asked whether T cells alone are sufficient to support VINCIT-mediated tumor control and in vivo CAR T cell generation. To address this, humanized NSG mice were reconstituted with defined PBMC subsets prior to tumor challenge and VINCIT treatment: purified T cells alone, T cells plus B cells, T cells plus monocytes, or unfractionated PBMCs (**Fig. 5a**). Tumor progression was monitored longitudinally by IVIS imaging (Fig. 5b–d).

**Figure 5.**
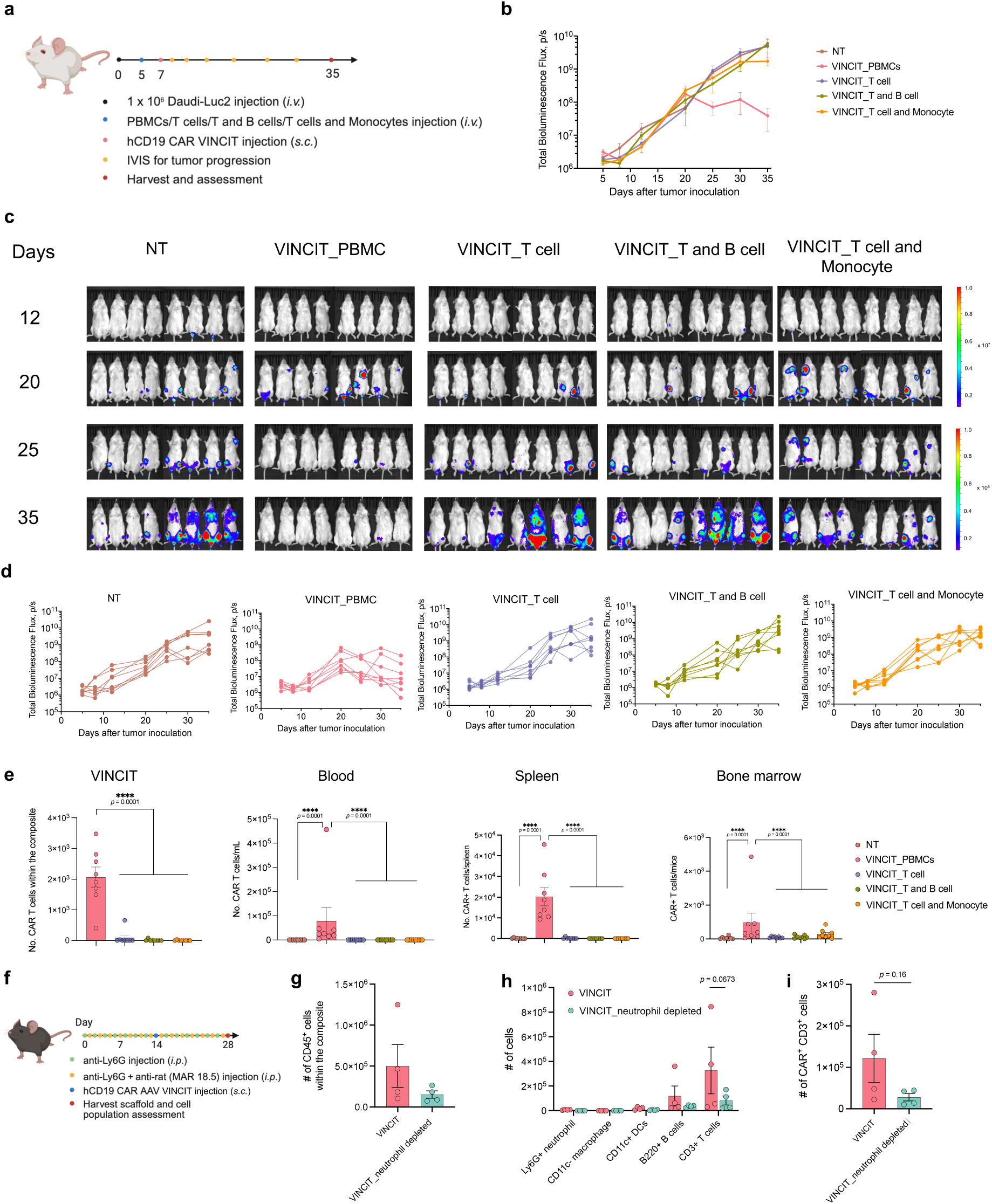
VINCIT orchestrates innate and adaptive immune cooperation to drive *in vivo* CAR T cell generation. **a**–**e,** Contribution of immune cell populations to VINCIT-mediated tumor control. **a,** Experimental timeline. NSG mice received intravenous injection of Daudi-Luc2 cells on day 0, followed by intravenous injection of defined cell populations isolated from PBMCs (T cells, or T cells + B cells, or T cells + B cells + monocytes) or complete PBMCs on day 5. hCD19 CAR AAV VINCIT was injected subcutaneously on day 7. Tumor progression was monitored by IVIS imaging. **b,** Representative bioluminescence images at the indicated time points. **c,** Average bioluminescence flux over time. **d**, Individual tumor burden curves for each group. **e,** Number of CAR T cells in VINCIT, blood, spleen, and bone marrow on day 35. **f**–**i,** Effect of neutrophil depletion on VINCIT function. **f**, Experimental timeline. Mice received anti-Ly6G antibody alone or anti-Ly6G plus anti-rat (MAR 18.5) antibody intraperitoneally every other day throughout the study. hCD19 CAR AAV VINCIT was injected subcutaneously on day 7. Scaffolds were harvested on day 28 for analysis. **g,** Total number of CD45^+^ cells within VINCIT. **h,** Quantification of immune cell populations within VINCIT, including Ly6G^+^ neutrophils, CD11c⁻ macrophages, CD11c^+^ dendritic cells, B220^+^ B cells, and CD3^+^ T cells. **i,** Percentage of CAR T cells among total live cells within VINCIT. Data are from n = 8 (a-e) or n = 4 (f-i) biologically independent samples. Data were analyzed using one-way ANOVA and Tukey’s multiple comparisons test. Panels **a** and **f** were created in BioRender.

Notably, all reconstitution groups recruited comparable total cell numbers within the VINCIT scaffold (**Supplementary Fig. 13**), suggesting that differences in tumor control were not attributable to differences in overall cellular infiltration. However, only mice reconstituted with complete PBMCs exhibited robust recruitment of human T cells to the scaffold (**Supplementary Figs. 14** and **15**), indicating that non-T cell populations play an active role in promoting T cell homing to, or expansion within, the VINCIT microenvironment. Consistent with this, *in vivo* CAR T cell generation within VINCIT was markedly reduced in mice lacking complete PBMC reconstitution (**Fig. 5d**), demonstrating that efficient CAR T cell generation depends on cooperative interactions among multiple immune cell populations. Furthermore, histopathological analysis of major organs four weeks after subcutaneous VINCIT administration revealed no overt histopathological abnormalities in the examined organs (**Supplementary Fig. 16**).

In addition, mice reconstituted with complete PBMCs displayed abundant CAR T cells within the VINCIT scaffold as well as in circulation, spleen, and bone marrow (**Fig. 5e**). In contrast, mice receiving T cells alone exhibited minimal CAR T cells across all compartments, consistent with impaired recruitment and/or expansion in the absence of supporting immune populations.

Phenotypic analysis revealed that VINCIT-generated CAR T cells in circulation and tissues were predominantly effector in phenotype, accompanied by detectable central memory and stem-like memory subsets (**Supplementary Fig. 17**), with low frequencies of PD-1⁺LAG-3⁺ exhausted cells across compartments (**Supplementary Fig. 18**). This phenotype is consistent with functional activation alongside the potential for sustained antitumor activity^33^.

We next investigated the role of innate immune cells in establishing this permissive immune niche. Because neutrophils are among the earliest responders to injected biomaterials, we investigated whether they help establish the immune microenvironment that supports subsequent cellular recruitment. To test this, we employed a dual–antibody-mediated depletion strategy to ablate neutrophils throughout the course of VINCIT treatment (**Fig. 5f**). Neutrophil depletion resulted in a pronounced reduction in total cellular infiltration within the VINCIT scaffold (**Fig. 5g**), affecting multiple downstream immune populations, including macrophages, dendritic cells, B cells, and CD3⁺ T cells (**Fig. 5h**). Consistent with this, *in vivo* CAR T cell generation within VINCIT was profoundly impaired (**Fig. 5i**), supporting an important early role for neutrophils in establishing the immune environment required for efficient CAR T cell generation. Together, these findings support a model in which VINCIT functions as a localized immune niche that coordinates innate and adaptive immune cell interactions to enable efficient *in vivo* CAR T cell generation. Rather than serving solely as a passive viral vector reservoir, VINCIT establishes a permissive microenvironment in which multicellular immune dynamics contribute to T cell recruitment, activation, and genetic programming.

## DISCUSSION

In this study, we developed VINCIT as an injectable biomaterial niche that couples spatially confined viral gene delivery with endogenous immune cell recruitment to enable CAR T cell generation in vivo. Importantly, localization of viral delivery alone was insufficient for efficient T cell programming: although free s.c. AAV produced substantial local transgene expression, it resulted in limited T cell transduction. By integrating viral retention with the immune cell–recruiting properties of the NHC, VINCIT markedly enhanced in situ T cell transduction and generated functional CAR T cells that entered the circulation and mediated systemic therapeutic activity. These findings establish the coupling of localized gene delivery and biomaterial-mediated immune cell recruitment as a key design principle for in vivo immune cell engineering.

VINCIT differs from existing approaches to simplifying CAR T cell generation in two important ways. Systemic in vivo delivery strategies bypass ex vivo cell processing but require vectors to reach and selectively transduce target T cells following whole-body administration, creating challenges associated with biodistribution, off-target exposure, and vector dose. VINCIT instead concentrates vector activity within a defined tissue compartment enriched with recruited immune cells, thereby separating the site of T cell programming from the eventual systemic distribution of the engineered cells. Conversely, previously reported biomaterial-based CAR T platforms provide a localized environment for T cell activation or transduction but generally require ex vivo isolation, engineering, or loading of T cells before implantation. For example, MASTER is preloaded with patient PBMCs and retroviral vector before subcutaneous implantation^23^, whereas VINCIT is injected without preloaded cells and recruits host T cells for transduction in situ. VINCIT combines the principal advantages of these approaches by recruiting endogenous cells directly into a localized transduction niche, enabling CAR T cell generation without an ex vivo cell-handling step.

The effectiveness of VINCIT appears to depend not only on recruitment of T cells, but on the establishment of a multicellular immune microenvironment that supports their subsequent programming. In humanized mice, T cells alone were insufficient to support robust CAR T cell generation, whereas reconstitution with complete PBMC populations markedly increased T cell recruitment and CAR T cell production within VINCIT. These findings suggest that non-T cell populations contribute to creating a permissive niche for T cell recruitment, activation, and/or expansion. Neutrophils appear to play an important early role in this process: their depletion reduced overall cellular infiltration and the recruitment of multiple downstream immune populations^34^, including T cells, and markedly impaired CAR T cell generation. Together, these observations support a model in which VINCIT functions as an immune-interactive niche, with early innate responses helping establish the multicellular environment required for efficient in situ T cell programming. Defining the molecular signals and cellular interactions underlying this process will be important for the rational design of future immune-engineering niches.

The ability to shift CAR T cell programming from an ex vivo manufacturing process to a localized in situ generation site has important translational implications. VINCIT does not require isolation, genetic modification, expansion, or reinfusion of T cells before administration, thereby consolidating multiple cell-processing steps into an injectable biomaterial–vector formulation. Spatial confinement may provide an additional advantage for viral gene delivery by concentrating vector exposure at a site enriched with recruited immune cells while limiting dissemination to non-target tissues. Importantly, the resulting engineered T cells are not restricted to the site of gene delivery but enter the circulation and traffic to systemic compartments, enabling a localized intervention to generate a systemic cellular response. Translating this concept will require optimizing vector tropism and dose, characterizing long-term gene-transfer and immunological responses, and validating it in models that more closely approximate human immune physiology.

Several limitations define important next steps for VINCIT. First, the viral vectors used here were not engineered for T cell-selective transduction. AAV and LVV were also used with different CAR constructs and readouts, so these experiments establish compatibility with both vector systems rather than their relative efficacy. Although spatial confinement limits vector dissemination, multiple immune populations enter the VINCIT niche and may therefore be exposed to the incorporated vectors. Combining localized delivery with T cell-targeted viral vectors could further improve cellular specificity and reduce unintended gene transfer. Second, the durability of genetic modification requires additional evaluation. Second, CAR expression was not tracked longitudinally. CAR T cells were detectable in blood, spleen, and bone marrow at endpoint, but defining persistence kinetics will require serial measurement of circulating CAR T cells and vector copy number. This is particularly important for integrating vectors such as LVV, for which insertional mutagenesis and long-term genomic safety remain important considerations^35,36,37^. Third, VINCIT relies on endogenous immune cell recruitment and multicellular interactions, which may vary with immune status, age, disease, and prior treatment^38^. Finally, the humanized mouse model used here does not fully reproduce human immune physiology or the pharmacological and anatomical constraints that will emerge at larger scale^39,40,41^. NSG mice retain myeloid cells but with impaired macrophage and dendritic cell function, biomaterial responses in this background are blunted ref:), and transferred human leukocytes act alongside residual murine populations. The reconstitution experiments therefore establish that non-T cell populations are required without quantifying each contribution, whereas the neutrophil requirement was defined in immunocompetent C57BL/6 mice. Human monocytes, macrophages, and NK cells were still detectable within VINCIT at day 35 (Supplementary Fig. 14), but a single time point cannot define subset persistence, and this model provides no human leukopoiesis to replenish short-lived populations; evaluating VINCIT in a model supporting human myeloid development is an important next step. Addressing these questions will require systematic evaluation of vector specificity, dose, persistence, and long-term responses in more clinically representative models.

Together, these findings establish VINCIT as a localized biomaterial niche for in vivo immune cell engineering. By coupling immune cell recruitment with spatially confined gene delivery, VINCIT enables therapeutic cells to be genetically programmed within a defined tissue compartment and subsequently deploy systemically. This strategy provides a framework for using engineered biomaterial niches to control where cellular programming occurs in vivo while preserving the capacity of the resulting cells to act beyond the site of intervention.

## METHODS

### Ethics statement

All animal procedures were performed under an animal protocol approved by the Johns Hopkins Institutional Animal Care and Use Committee (protocol #MO24E431). Male and female C57BL/6 mice, 6–8 weeks of age, were purchased from the Jackson Laboratory. Male and female NSG mice, 6–8 weeks of age, were bred in Johns Hopkins Animal Care Facilities and randomly grouped. Mice had unlimited access to pelleted food and water throughout the study. The maximum tumor size permitted by the IACUC was 1 × 10^9^ p/s or a weight loss of more than 15%. Mice were euthanized when either threshold was reached. We confirm that these limits were not exceeded in any of the experiments reported in this study.

### Chemicals and reagents

Sodium hyaluronate (HA, molecular weight 1.01–1.80 MDa, research grade) was purchased from LifeCore Biomedical Inc (Chaska, MN, USA). Glycidyl acrylate was obtained from TCI America Inc. (Portland, OR, USA). Poly(ε-caprolactone) (PCL, molecular weight 80 kDa), glycidyl acrylate (GA), and all other chemical reagents were purchased from Sigma-Aldrich unless otherwise noted.

### Viral vectors

AAV9-CMV-tdTomato and AAV9-CMV-luciferase were obtained from Virovek. AAV6 encoding human CD19 CAR (CD19 scFv-CD8^-^4-1BB-CD3ζ under EF1α promoter) was obtained from BPS Bioscience. Lentiviral vector encoding mouse CD19 CAR (scFv-CD8 hinge-m4-1BB-mCD3ζ under EF1α promoter with CopGFP reporter) was obtained from Creative Biolabs. LVV-CMV-tdTomato was obtained from Porton Advanced Solutions. AAV was labeled with Cy7.5 by reacting Cy7.5-NHS ester with primary amines on AAV capsid proteins according to standard NHS ester conjugation protocols.

### Preparation of VINCIT

NHC was synthesized as previously described. Briefly, HA was acrylate-modified by reacting with GA in phosphate-buffered saline (PBS, pH 8.5) at 37°C for 16 h. PCL nanofibers were electrospun from a 16% (w/w) PCL solution in 9:1 (v/v) dichloromethane and dimethylformamide through a 27-G needle at 2.5 mL/h with 16 kV electric potential. Electrospun fibers were plasma-treated to generate carboxylic groups, activated with EDC/NHS, and functionalized with maleimide groups. Fibers were cryogenically milled to generate fragments with an average length of approximately 50 μm. NHC was prepared by mixing maleimide-modified PCL fiber fragments (30 mg/mL) with HA-Ac solution (10 mg/mL in PBS, pH 7.4) and PEG-SH (5 mg/mL). Crosslinking was mediated by Michael addition reaction at 37°C. The NHC was mechanically fragmented by passing through a stainless-steel mesh with 50 μm pore opening to generate microgels. ViHA (hydrogel alone) was prepared using the same protocol without PCL fibers. Viral vectors were incorporated into NHC by mixing at a 1:1 volume ratio using two syringes connected with a connector. The mixture was used immediately for injection. AAV was used at 10^10^ vg per injection and lentiviral vectors at 10^7^ TU per injection.

### Cells

Daudi-Luc2 cells (ATCC CCL-213-LUC2), a human Burkitt’s lymphoma B cell line stably expressing luciferase, were cultured in RPMI-1640 medium supplemented with 10% fetal bovine serum (FBS) and 4 µg/mL Blasticidin, and maintained at 37°C in a humidified incubator with 5% CO_2_.

Peripheral blood mononuclear cells (PBMCs) were isolated from leukopaks obtained from anonymous healthy donors. Briefly, leukopak contents were diluted with phosphate-buffered saline (PBS) containing 2% FBS and carefully layered over Ficoll-Paque density gradient medium (Cytiva) in conical tubes. Samples were centrifuged at 400 × g for 30 min at room temperature with the brake off. The PBMC layer at the interface was carefully collected and washed twice with PBS containing 2% FBS. Cells were counted using a hemocytometer or automated cell counter and either used immediately for downstream applications or cryopreserved in FBS containing 10% DMSO for future use.

T cells, B cells, and monocytes were isolated from human PBMCs using negative selection immunomagnetic kits (STEMCELL Technologies). T cells were isolated using the EasySep Human T Cell Isolation Kit (Cat #17951), B cells using the EasySep Human B Cell Isolation Kit (Cat #17954), and monocytes using the EasySep Human Monocyte Isolation Kit (Cat #19359). Briefly, PBMCs were resuspended at the recommended concentration and incubated with the isolation cocktail containing antibodies targeting non-desired cell types. Following incubation, magnetic particles were added, and samples were placed in an EasySep magnet. Untouched, highly purified target cells were collected by pouring off the enriched fraction while magnetically labeled unwanted cells remained bound inside the tube. Isolation was performed according to the manufacturer’s instructions.

### Animals

C57BL/6J mice and NOD.Cg-Prkdcscid Il2rgtm1Wjl/SzJ (NSG) mice were purchased from The Jackson Laboratory. Mice were 6–8 weeks old at the start of experiments, and both male and female mice were used in equal proportions. All animal procedures were approved by the Johns Hopkins University Institutional Animal Care and Use Committee. The feed composition consisted of approximately 5% fiber, 20% protein, and 5–10% fat. Daily food consumption averaged 4–5 g per mouse (roughly 120 g/kg body weight), while water intake ranged from 3–5 mL daily (approximately 150 mL/kg body weight). Housing facilities were kept at 18–26°C (64–79°F) with relative humidity between 30% and 70%, and ventilation provided at least 10 air exchanges per hour. Mice were housed in standard shoebox cages with corncob bedding material. The AAV or VINCIT were given through s.c. (right flank) injection at a predetermined dose per mouse.

### Biodistribution studies

C57BL/6 mice received Cy7.5-labeled AAV (10^10^ vg in 200 μL) via intravenous injection, or subcutaneous injection in free form or VINCIT. At days 15 and 30, mice were euthanized and organs harvested for imaging. For bioluminescence studies, mice received luciferase-encoding AAV and were imaged using the Xenogen IVIS Imaging System with Living Image 4.2 software. Total flux was quantified from regions of interest.

### Scaffold harvest and cell isolation

At the indicated time points, scaffolds were surgically excised, with the surrounding tissue carefully removed. Scaffolds were digested with hyaluronidase (7 mg/mL) at 37°C for 30 min. Cell suspensions were passed through 40-μm cell strainers and pelleted by centrifugation at 300 × g for 5 min. Red blood cells were lysed with ACK lysis buffer when necessary.

### Flow cytometry

Cells were stained with antibodies in flow cytometry staining buffer for 20 min at 4°C in the dark. Antibodies used in the study are Live/Dead Fixable Aqua (Invitrogen # L34966), Zombie Yellow Fixable Viability Kit (Biolegend # 423104); BV421 anti-mouse CD45 (Biolegend # 103134), APC anti-mouse CD3 (Biolegend # 100236), PE-Cy7 anti-mouse CD45R (B220) (Biolegend # 103221), BV785 anti-mouse CD11b (Biolegend #101243), APC-Cy7 anti-mouse CD11c (Biolegend # 117324), PerCP-Cy5.5 anti-mouse Ly6G (Biolegend #127616), FITC anti-mouse MHC II (Biolegend # 125508).

BV421 anti-human CD45 (Biolegend #368522), APC, PE anti-human CD3 (Biolegend # 317318, 317308), FITC anti-human CD4 (Biolegend #317408), BV510 anti-human CD8a (Biolegend # 301048), BV650 anti-human CD19 (Biolegend #302238), APC-Cy7 anti-human CD45RA (Biolegend # 304128), APC anti-human CCR7 (Biolegend #353214), PerCP-Cy5.5 anti-human PD-1 (Biolegend # 367410), PE-Cy7 anti-human LAG-3 (Biolegend #369310). PE-Labeled Monoclonal Anti-FMC63 (Acrobiosystems, FM3-HPY53). All antibodies were used at 1:100 dilution. Data were acquired on an Attune NxT flow cytometer and analyzed using FlowJo v11 software.

### Intracellular cytokine staining

Splenocytes were stimulated with Daudi cell lysate (2 μg protein/mL) for 18 h in the presence of brefeldin A. Cells were stained with Live/Dead Fixable Aqua (Invitrogen # L34966), APC anti-mouse CD3 (Biolegend # 100236), APC-Cy7 anti-mouse CD8 (Biolegend # 100714), then fixed, permeabilized, and stained with PE-Cy7 anti-mouse Granzyme B (Biolegend #396410), AF700 anti-mouse TNF-α (Biolegend #506338), and BV650 anti-mouse IFN-γ (BD #563854).

### FluoroSpot assay

IFN-γ production by mouse T cells was assessed using the FluoroSpot Plus Mouse IFN-γ kit (Mabtech, Cat# FSP-4124-10). Briefly, PVDF membrane plates were pre-wetted with 35% ethanol and washed with sterile PBS. Plates were then coated with the provided anti-mouse IFN-γ capture antibody (AN18) and incubated overnight at 4°C. Following washing with PBS, plates were blocked with culture medium for at least 30 min at room temperature. Cells were seeded at the desired density and stimulated with the appropriate antigen or control for 24–48 h at 37°C with 5% CO_2_. After incubation, cells were removed, and plates were washed with PBS containing 0.1% Tween-20. Detection antibody (R4-6A2-490) was added and incubated for 2 h at room temperature. Following additional washes, the fluorescence enhancer was added and incubated for 15 min. Plates were emptied, dried, and spots were visualized and counted using an automated FluoroSpot reader.

### Serum cytokine analysis

Serum was collected at day 3 post-injection. IL-6, TNF-α, IL-10, IL-12, and IFN-γ were measured using ELISA kits (ThermoFisher # 88-7064-88, 88-7324-88, 88-7105-88, 88-7121-88, 88-7316-88) according to manufacturer’s protocols.

### B cell depletion studies

C57BL/6 mice received mCD19 CAR LVV via VINCIT subcutaneously or intravenous injection. Peripheral blood was collected via submandibular bleeding at days 10 and 20. Blood was processed with ACK lysis buffer and stained for CD45, CD3, and CD19 for flow cytometry analysis.

### Humanized mouse tumor model

NSG mice received 1 × 10^6^ luciferase-expressing Daudi cells (ATCC #CCL-213-LUC2) intravenously on day 0. On day 5, mice received 1 × 10^7^ human PBMCs intravenously. On day 7, mice were treated with hCD19 CAR AAV via VINCIT subcutaneously, intravenous injection, or controls. Tumor burden was monitored by bioluminescence imaging on days 7, 11, 14, 25, 32, 40, 54, and 60. Mice were euthanized when weight loss exceeded 15% or bioluminescence exceeded 1 × 10^9^ photons/second.

### Neutrophil depletion

Neutrophils were depleted as previously described. Mice received anti-mouse Ly6G antibody (clone 1A8, Bio X Cell, 200 μg per injection) and anti-rat kappa immunoglobulin light chain antibody (clone MAR 18.5, Bio X Cell, 100 μg per injection) intraperitoneally every other day throughout the study period.

### Immune cell population isolation

T cells, B cells, and monocytes were isolated from human PBMCs using magnetic bead isolation kits (STEMCELL Technologies) according to the manufacturer’s protocols. Purity of isolated populations exceeded 95% as confirmed by flow cytometry.

### Scanning electron microscopy

VINCIT scaffolds were lyophilized and cut with a sharp razor blade to expose cross-sections. Samples were sputter-coated with gold-palladium and imaged using a scanning electron microscope (Thermo Fisher Scientific Helios G4 UC, USA) at an accelerating voltage of 10 kV. Images were used to characterize scaffold morphology, pore structure, and fiber distribution. Pore size distribution was quantified using ImageJ Fiji (NIH, Bethesda, MD) (**Supplementary Fig. 1b**).

### Histological analysis

C57BL/6J mice were injected subcutaneously with VINCIT loaded with empty AAV (n = 3 per group). Untreated mice served as controls. Mice were euthanized four weeks after injection. Major organs (heart, liver, kidney, lung, and spleen) and the injection site were excised and fixed in 10% neutral buffered formalin for 24 h at room temperature. Fixed tissues were processed, embedded in paraffin, sectioned at 5 μm thickness, and stained with hematoxylin and eosin (H&E). Slides were scanned using a NanoZoomer digital slide scanner (Hamamatsu Photonics) and assessed for tissue morphology and histopathological evidence of inflammation or injury (**Supplementary Fig. 18**).

### Statistical analysis

Statistical analyses were performed using GraphPad Prism. Comparisons between two groups were made using an unpaired two-tailed Student’s t-test. Comparisons among multiple groups were made using one-way or two-way ANOVA with Tukey’s post hoc test. Survival curves were analyzed using the log-rank (Mantel-Cox) test. Data are presented as mean ± s.e.m. unless otherwise indicated.

## Data availability

All data generated or analysed during this study are provided in the paper and the Supplementary Information. Source data supporting the findings of this study are available in the accompanying Source Data file. No additional datasets requiring deposition in external repositories were generated. Source data are provided in this paper.

## Acknowledgements

This study was partially supported by the National Institutes of Health/National Cancer Institute through grant R01 CA293906 (H.-Q.M. and J.P.S.), the National Institute of Biomedical Imaging and Bioengineering through grant P41 EB028239 (J.P.S. and H.-Q.M.), and the National Institute of Allergy and Infectious Diseases through grant R21 AI176764 (M.F.K.).

## Author Contributions

K.D.G., Y.Z., and H.Q.M. conceptualized and designed this study. H.Q.M., J.P.S., and M.F.K. secured the funding for this study. K.D.G., Y.Z., C.W., Z.J., Y.J., X. Lu, S.C., A.H.G., D.Y., X. Liu, J.C., Y.S., J.M., J.L., J.K., E.W., and D.L. performed the experiments. D.S.W., J.C.D., J.P.S., and M.F.K. participated in data analysis and interpretation. The manuscript was written by K.D.G., Y.Z., and H.Q.M., with revisions by D.S.W., J.C.D., J.P.S., and M.F.K, and input from all the other authors. All authors approved the submitted manuscript version.

## Competing Interests

H.-Q.M., K.D.G. and Y.Z. are co-inventors of a patent application covering the VINCIT platform described in this paper, filed through and managed by Johns Hopkins Technology Ventures. The other authors declare no competing interests.

