## Supplemental Information for "VINCIT: A virus-incorporated biomaterial niche enables spatially controlled in vivo CAR T cell engineering"

### Supplementary Figures

**Figure 1.** Rheological characterization of VINCIT.

**Figure 2.** VINCIT injection and morphology characterization.

**Figure 3.** Ex vivo fluorescence imaging of harvested organs following Cy7.5-labeled AAV delivery.

**Figure 4.** Organ-level quantification of Cy7.5-labeled AAV biodistribution.

**Figure 5.** Bioluminescence imaging of harvested organs following Luc AAV delivery at day 30.

**Figure 6.** Longitudinal bioluminescence signal at VINCIT injection sites.

**Figure 7.** Transduction efficiency and T cell specificity at the injection site.

**Figure 8.** Flow cytometry gating strategy for characterization of immune cell recruitment and transduction within VINCIT scaffolds.

**Figure 9.** Individual tumor burden for NT and VINCIT_PBMC groups.

**Figure 10.** Systemic cytokine levels following VINCIT injection.

**Figure 11.** Flow cytometry gating strategy for identification of in vivo-generated CAR-T cells.

**Figure 12.** Flow cytometry gating strategy for phenotypic characterization of VINCIT-generated CAR-T cells.

**Figure 13.** Cell recruitment within VINCIT across immune cell composition groups.

**Figure 14.** Immune cell composition within VINCIT in the PBMC group at endpoint.

**Figure 15.** T cell percentage of $CAR^+$ cells across tissue compartments.

**Figure 16.** Representative images of H&E-stained sections of five major organs four weeks after subcutaneous injection of VINCIT with empty AAV and untreated controls in C57Bl6/J mice.

**Figure 17.** Phenotypic characterization of VINCIT-generated CAR-T cells humanized with PBMCs.

**Figure 18.** Exhaustion profile of VINCIT-generated CAR-T cells humanized with PBMCs.

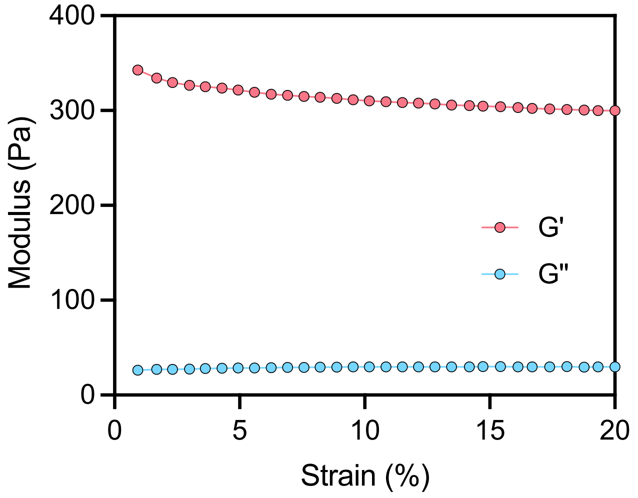


**Supplementary Figure 1. Rheological characterization of VINCIT.** Representative storage modulus (G') and loss modulus (G") as a function of strain (0–20%).

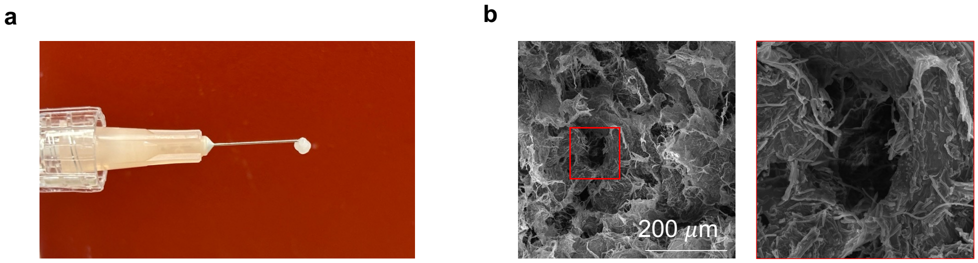


**Supplementary Figure 2. VINCIT injection and morphology characterization. a**, Photograph demonstrating injectability of VINCIT through a 30-gauge needle. **b**, Scanning electron microscopy (SEM) images of VINCIT showing macroporous structure. Left: low magnification view. Right: magnified view of the boxed region showing interconnected pores with an average pore size of approximately 50 μm. Scale bar, 200 μm.

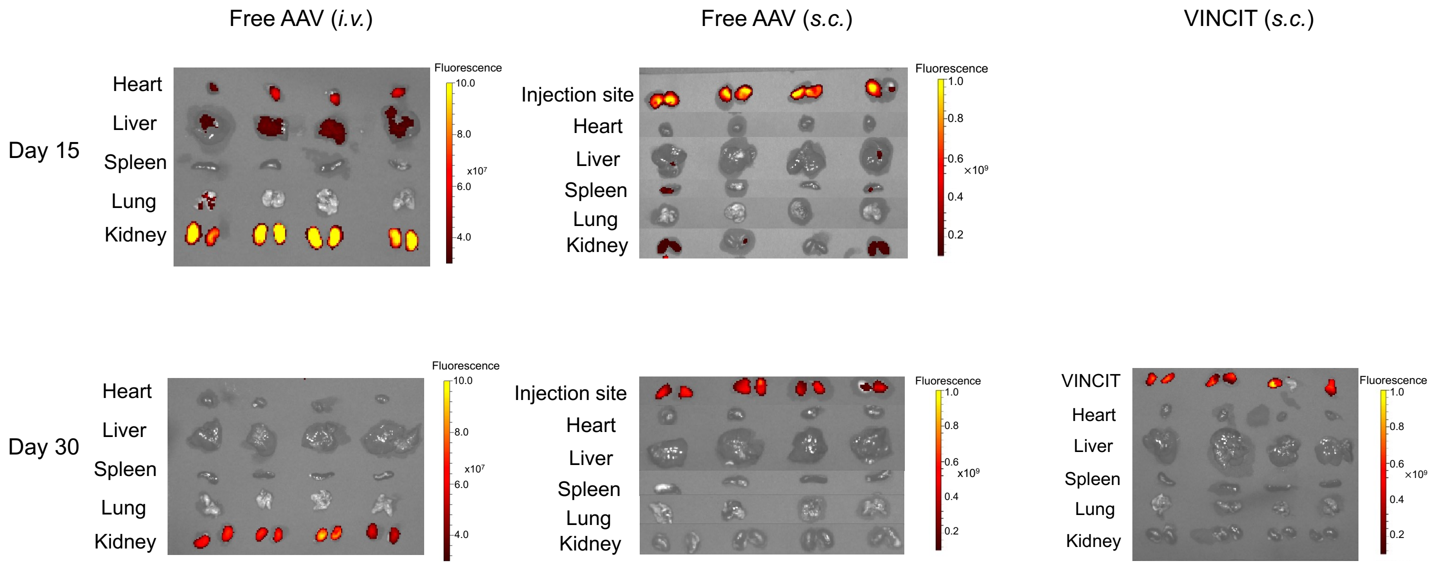


**Supplementary Figure 3. Ex vivo fluorescence imaging of harvested organs following Cy7.5-labeled AAV delivery.** Representative fluorescence images of major organs (heart, liver, spleen, lung, kidney) and VINCIT scaffolds harvested from mice at days 15 and 30 after intravenous injection (i.v.), subcutaneous injection (s.c.), or subcutaneous injection of AAV-loaded VINCIT. Fluorescence intensity is displayed as a pseudocolor scale ($1 \times 10^9$, radiant efficiency [p/s/cm²/sr]/[μW/cm²]). n = 4 biologically independent samples.

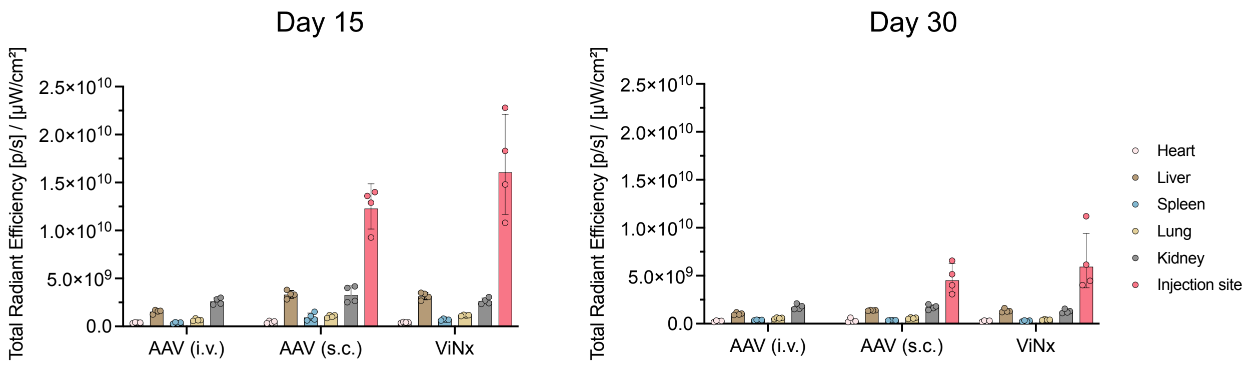


**Supplementary Figure 4. Organ-level quantification of Cy7.5-labeled AAV biodistribution.** Fluorescence intensity of individual organs harvested from mice at day 15 (left) and day 30 (right) following intravenous injection (i.v.), subcutaneous injection (s.c.), or subcutaneous injection of AAV-loaded VINCIT. n = 4 biologically independent samples. Data represent mean ± s.e.m.

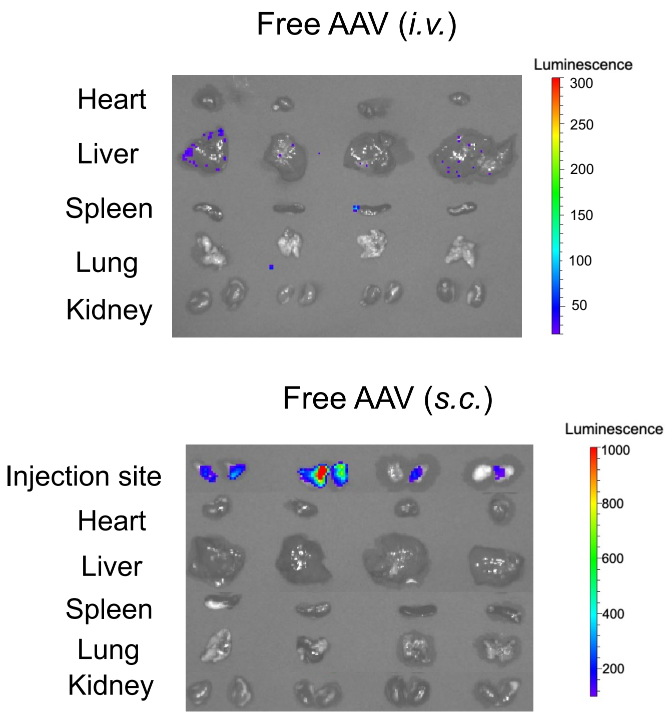


**Supplementary Figure 5. Bioluminescence imaging of harvested organs following Luc AAV delivery.** Representative ex vivo bioluminescence images of major organs (heart, liver, spleen, lung, kidney) harvested at day 30 from mice receiving Luc AAV via intravenous injection (i.v.), subcutaneous injection (s.c.). Luminescence intensity is displayed as a pseudocolor scale (photons/s/$cm^2$/sr). n = 4 biologically independent samples.

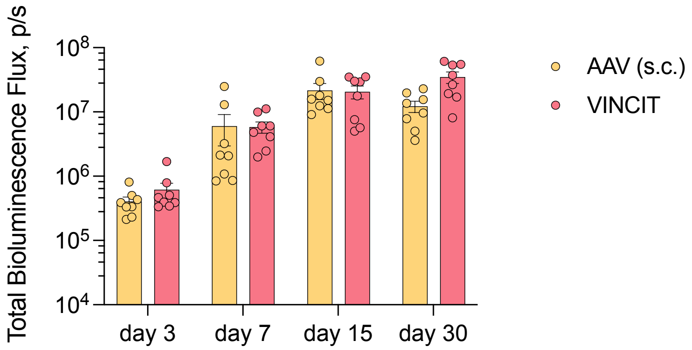


**Supplementary Figure 6. Longitudinal bioluminescence signal at VINCIT injection sites.** Quantification of luciferase expression at individual injection sites over 30 days following subcutaneous injection of Luc AAV-loaded VINCIT. Each mouse received bilateral flank injections. n = 4 mice (8 injection sites). Data represent mean ± s.e.m.

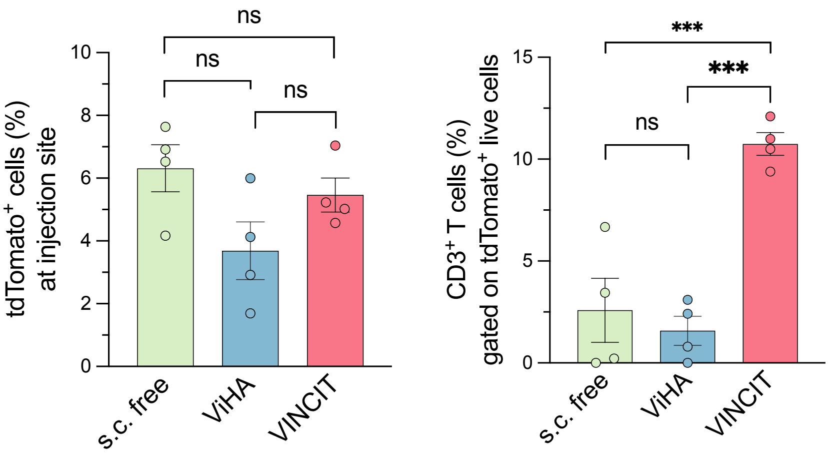


**Supplementary Figure 7. Transduction efficiency and T cell specificity at the injection site.** (a) tdTomato$^+$ cells (%) among live cells at the injection site. (b) CD3$^+$ T cells (%) gated on tdTomato$^+$ live cells. Mean ± SEM, n = 4 mice per group; one-way ANOVA with Tukey's test; ns, not significant; ***$p < 0.001$.

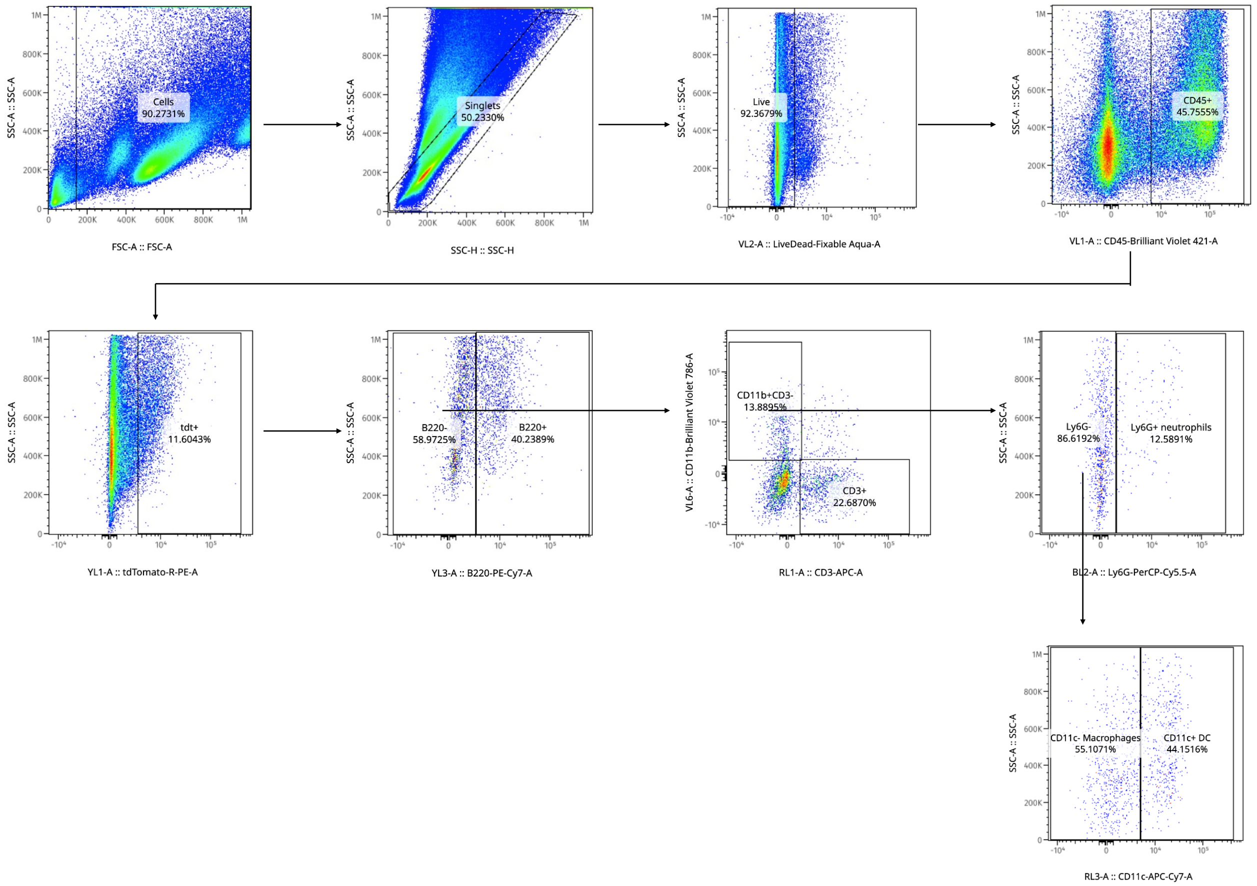


**Supplementary Figure 8. Flow cytometry gating strategy for characterization of immune cell recruitment and transduction within VINCIT scaffolds.** Representative flow cytometry plots depicting the sequential gating strategy used to identify host immune cell populations recruited to VINCIT scaffolds and assess viral transduction. $B220^+$ B cells, $B220^-$ $CD3^+$ T cells, $CD11b^+Ly6G^+$ neutrophils, $Ly6G^-$ $CD11c^-$ macrophages, and $Ly6G^-CD11c^+$ dendritic cells were gated.

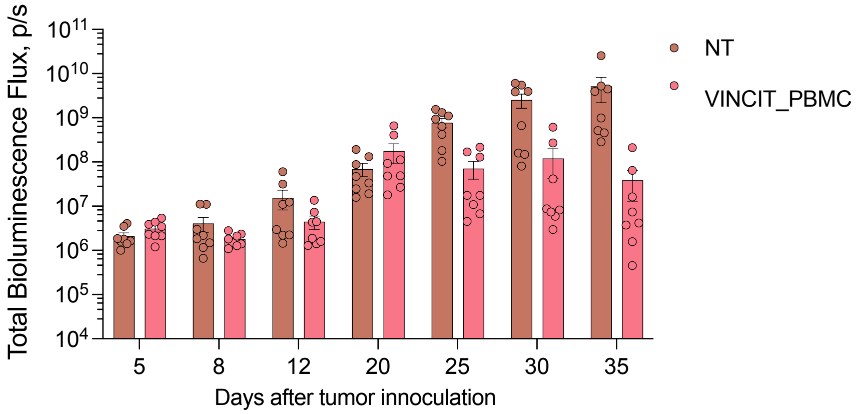


**Supplementary Figure 9. Individual tumor burden for NT and VINCIT_PBMC groups**. Bioluminescence flux for individual mice in the no treatment (NT) and VINCIT_PBMC groups at indicated time points, corresponding to the summary data in Fig. 5d. n = 8 biologically independent samples. Data represent mean ± s.e.m.

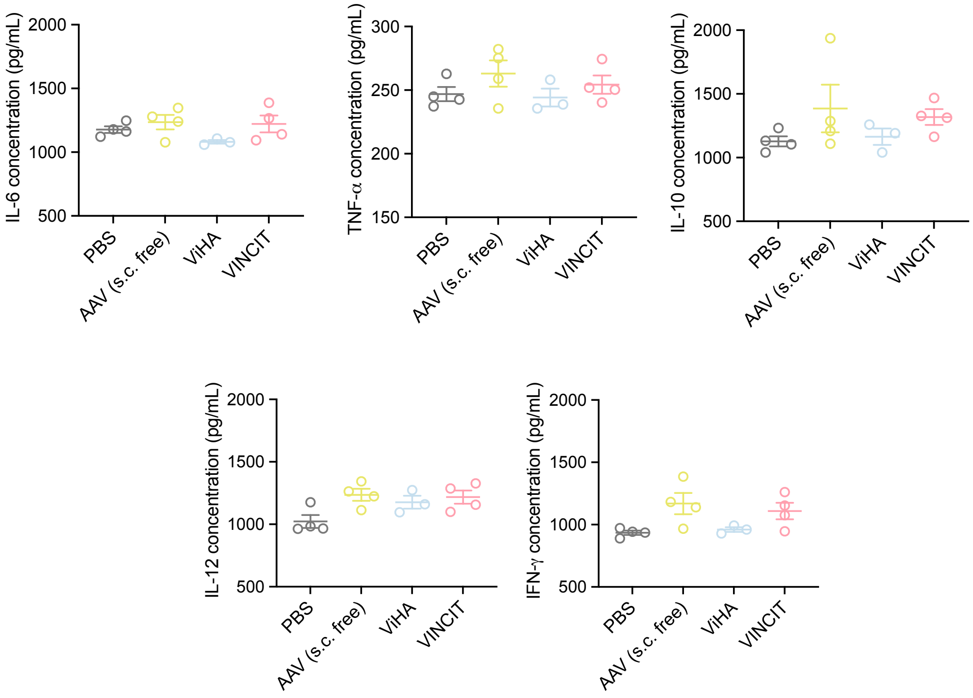


**Supplementary Figure 10. Systemic cytokine levels following VINCIT injection.** Serum cytokine concentrations at day 3 following subcutaneous injection of PBS, free AAV (s.c.), ViHA, or VINCIT loaded with empty AAV in C57BL/6 mice. IL-6, TNF-α, IL-10, IL-12, and IFN-γ were measured by ELISA. No significant differences were observed across groups. n = 4 biologically independent samples. Data represent mean ± s.e.m.

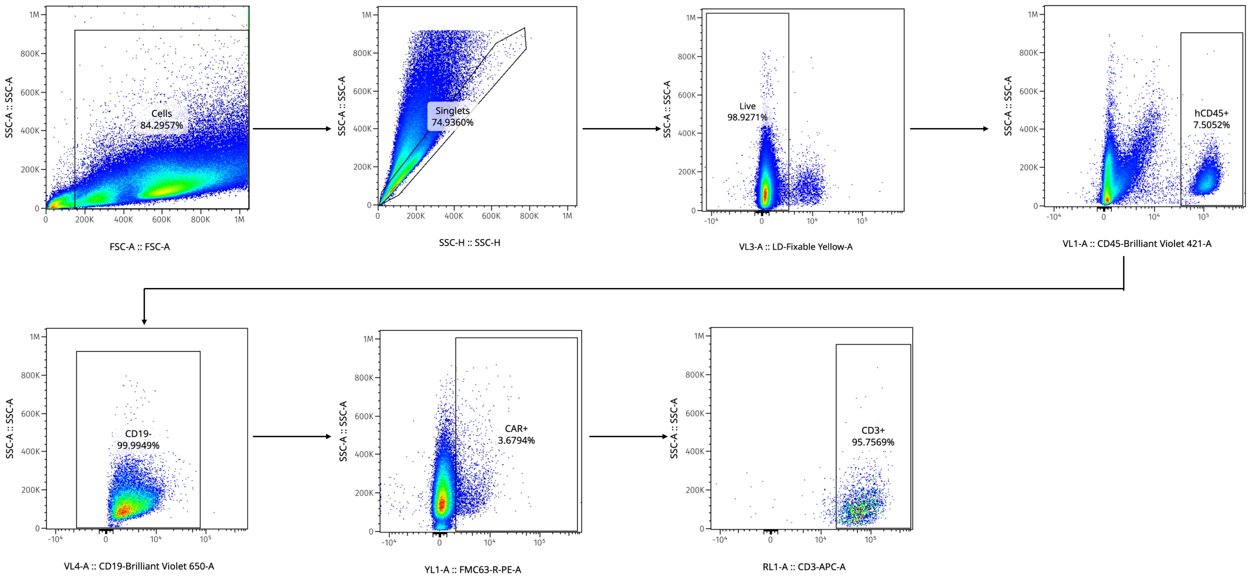


**Supplementary Figure 11. Flow cytometry gating strategy for identification of in vivo-generated CAR-T cells.** Representative flow cytometry plots depicting the sequential gating strategy used to identify CAR-T cells generated within VINCIT scaffolds. Cells were gated sequentially for intact cells, singlets, live cells, and hCD45$^+$ human leukocytes. B cells were excluded by gating on the hCD19$^-$ population, and CAR expression was detected using an FMC63-specific antibody. T cell identity was confirmed by CD3 staining, verifying that VINCIT-generated CAR$^+$ cells were of T cell lineage.

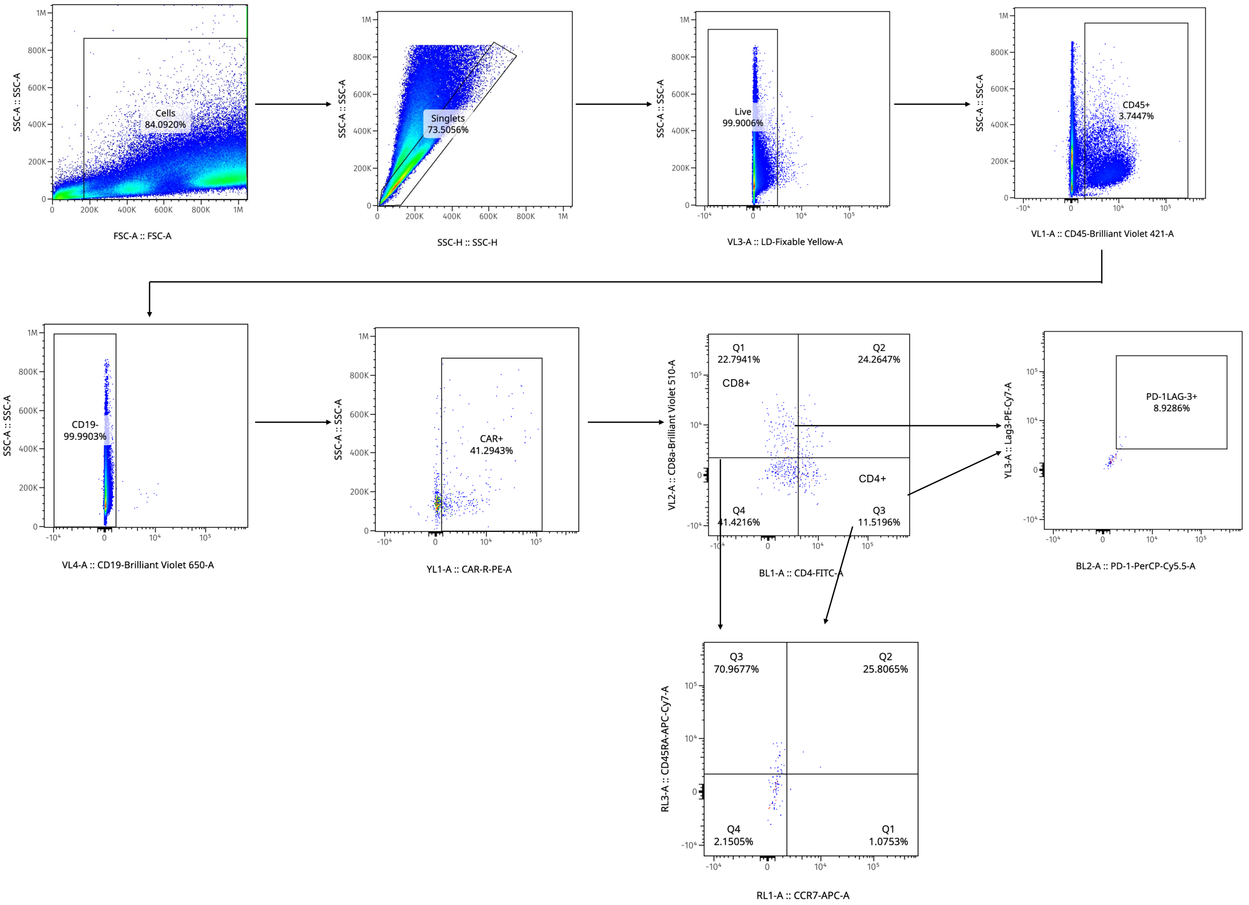


**Supplementary Figure 12. Flow cytometry gating strategy for phenotypic characterization of VINCIT-generated CAR-T cells.** Representative flow cytometry plots depicting the sequential gating strategy used to identify and characterize in vivo-generated CAR-T cells from VINCIT scaffolds. Cells were gated sequentially for intact cells, singlets, live cells, hCD45$^+$ leukocytes, and hCD19$^-$ non-B cells. CAR expression was assessed on the CD19$^-$ population. CAR$^+$ T cells were further characterized for CD4/CD8 subset distribution, exhaustion marker co-expression (PD-1/LAG-3), and memory differentiation status (CCR7/CD45RA).

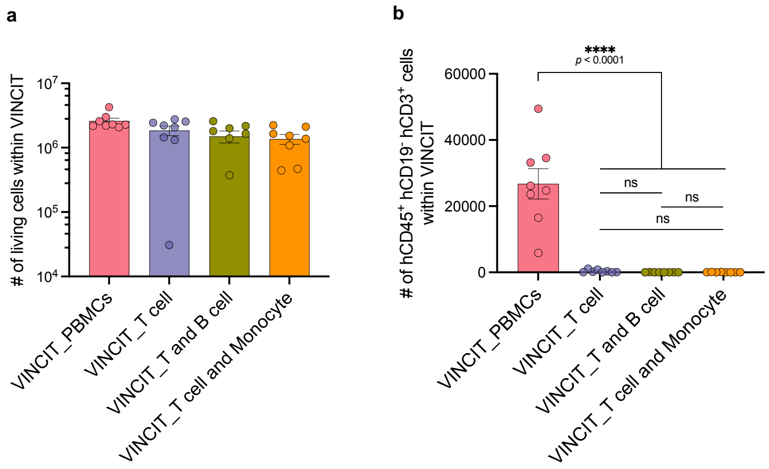


**Supplementary Figure 13. Cell recruitment within VINCIT across immune cell composition groups.** Total number of recruited living cells (**a**) and $hCD45^{+}hCD19^{-}hCD3^{+}$ T cells (**b**) within VINCIT scaffolds at endpoint for each immune cell composition group. Data represent mean ± s.e.m. Statistical comparisons were performed using one-way ANOVA with Tukey's multiple comparisons test. ns = no significance, ****$P < 0.0001$.

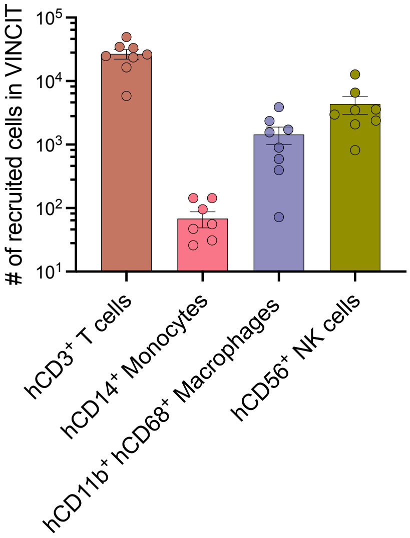


**Supplementary Figure 14. Immune cell composition within VINCIT in the PBMC group at endpoint.** Quantification of recruited hCD3$^+$ T cells, hCD14$^+$ monocytes, hCD11b$^+$hCD68$^+$ macrophages, and hCD56$^+$ NK cells within VINCIT scaffolds at endpoint for mice receiving PBMCs, corresponding to the tumor study in Fig. 5. n = 8 biologically independent samples. Data represent mean ± s.e.m.

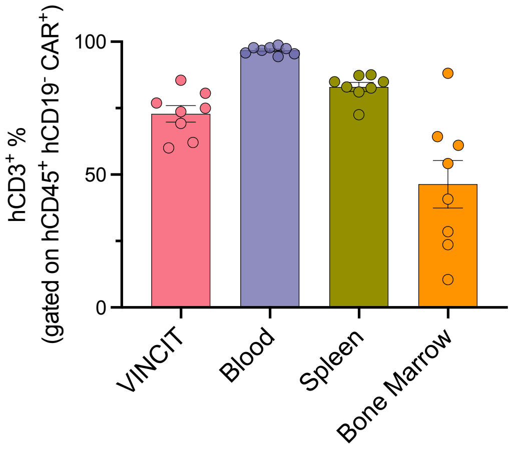


**Supplementary Figure 15. T cell percentage of $CAR^+$ cells across tissue compartments.** Percentage of $hCD3^+$ cells among $hCD45^+hCD19^-CAR^+$ cells in VINCIT, blood, spleen, and bone marrow at endpoint for the PBMC group. n = 8 biologically independent samples. Data represent mean ± s.e.m.

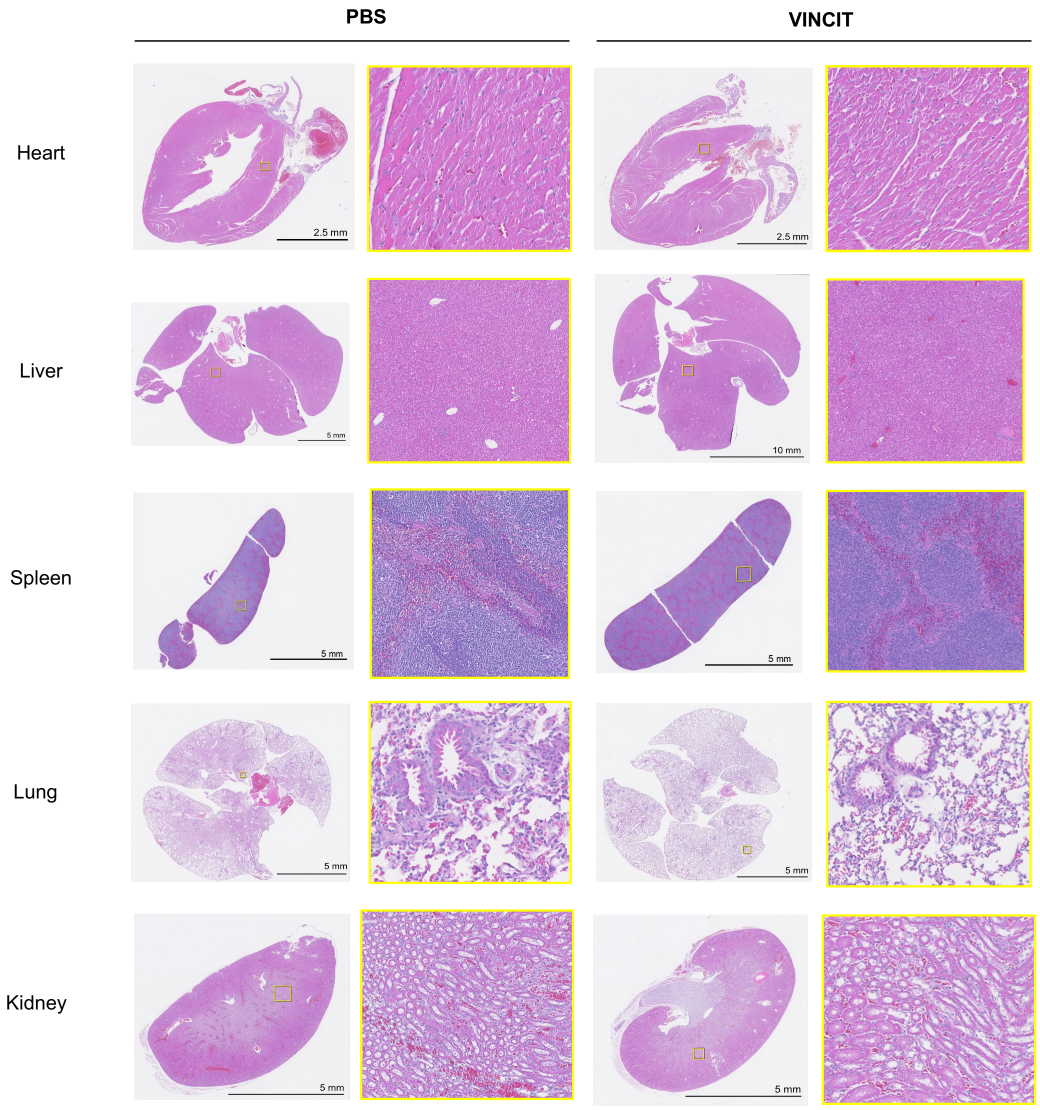


**Supplementary Figure 16. Representative images of H&E-stained sections of five major organs four weeks after subcutaneous injection of VINCIT with empty AAV and untreated controls in C57Bl6/J mice**. Data is representative of four biologically independent animals.

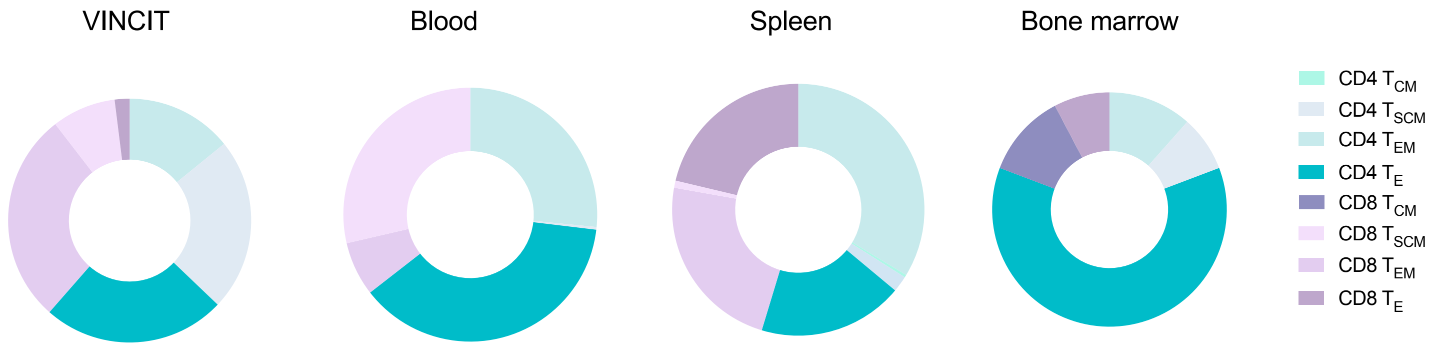


**Supplementary Figure 17. Phenotypic characterization of VINCIT-generated CAR-T cells humanized with PBMCs.** Memory phenotype of CAR-T cells isolated from VINCIT, blood, spleen, and bone marrow of humanized mice treated with hCD19 CAR AAV VINCIT. T cell subsets were defined by CD45RA and CCR7 expression: naïve/stem cell memory (TN/SCM): $CD45RA^{+}CCR7^{+}$; central memory (TCM): $CD45RA^{-}CCR7^{+}$; effector memory (TEM): $CD45RA^{-}CCR7^{-}$; effector (TE): $CD45RA^{+}CCR7^{-}$. Phenotypes are shown separately for $CD4^{+}$ and $CD8^{+}$ CAR-T cells.

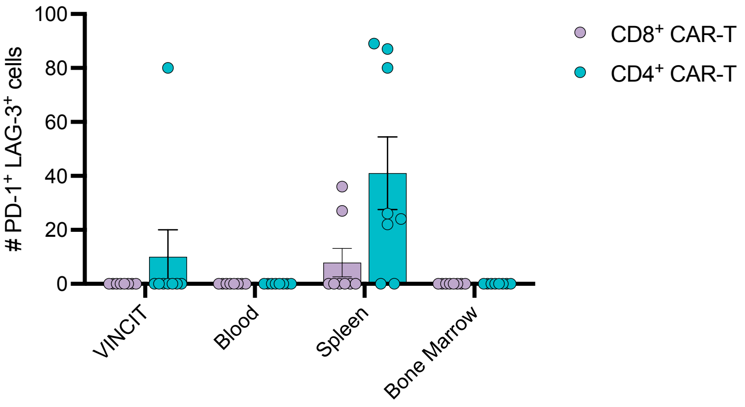


**Supplementary Figure 18. PD-1 and LAG-3 co-expression on VINCIT-generated CAR T cells humanized with PBMCs.** Number of PD-$1^+$LAG-$3^+$ CAR-T cells among $CD4^+$ and $CD8^+$ subsets in VINCIT, blood, spleen, and bone marrow of humanized mice treated with hCD19 CAR AAV VINCIT. n = 8 biologically independent samples. Data represent mean ± s.e.m.
